# Lipase-mediated detoxification of host-derived antimicrobial fatty acids by *Staphylococcus aureus*

**DOI:** 10.1101/2023.05.15.540481

**Authors:** Arnaud Kengmo Tchoupa, Ahmed M. A. Elsherbini, Xiaoqing Fu, Oumayma Ghaneme, Lea Seibert, Marieke A. Böcker, Marco Lebtig, Justine Camus, Stilianos Papadopoulos Lambidis, Birgit Schittek, Dorothee Kretschmer, Michael Lämmerhofer, Andreas Peschel

## Abstract

Long-chain fatty acids with antimicrobial properties are abundant on the skin and mucosal surfaces, where they are essential to restrict the proliferation of opportunistic pathogens such as *Staphylococcus aureus*. These antimicrobial fatty acids (AFAs) elicit bacterial adaptation strategies, which have yet to be fully elucidated. Characterizing the pervasive mechanisms used by *S. aureus* to resist AFAs could open new avenues to prevent pathogen colonization. Here, we identify the *S. aureus* lipase Lip2 as a novel resistance factor against AFAs. Lip2 detoxifies AFAs via esterification with cholesterol. This is reminiscent of the activity of the fatty acid-modifying enzyme (FAME), whose identity has remained elusive for over three decades. *In vitro*, Lip2-dependent AFA-detoxification was apparent during planktonic growth and biofilm formation. Our genomic analysis revealed that prophage-mediated inactivation of Lip2 was more common in blood and nose isolates than in skin strains, suggesting a particularly important role of Lip2 for skin colonization. Accordingly, in a mouse model of *S. aureus* skin colonization, bacteria were protected from sapienic acid - a human-specific AFA - in a cholesterol- and lipase-dependent manner. These results suggest Lip2 is the long-sought FAME that exquisitely manipulates environmental lipids to promote bacterial growth. Our data support a model in which *S. aureus* exploits and/or exacerbates lipid disorders to colonize otherwise inhospitable niches.

## Introduction

At the host-pathogen interface, lipids exert multifaceted functions as, for instance, building blocks for cells and extracellular matrices^1–3^, energy sources^4, 5^, entry routes into host cells^6^, immunomodulators^7^, and potent antimicrobials^8–10^. To harness environmental lipids and fuel their growth, bacteria utilize a plethora of lipolytic enzymes, whose substrates include sphingolipids, phospholipids, and triacylglycerols^4,11–13^. These lipid hydrolases release host-derived long-chain fatty acids with antibacterial properties, also referred to as antimicrobial fatty acids (AFAs)^14^. An intriguing concept is that bacteria would secrete lipases to release AFAs from complex lipids and thereby inhibit AFA-susceptible competitors within the same niche, for instance on human skin. This has been demonstrated for *Corynebacterium accolens* and *Streptococcus pneumoniae*^12^. Hence, adaptation strategies to AFAs represent a prerequisite for stable colonization of the skin and mucosal surfaces. *Staphylococcus aureus*, an opportunistic pathogen colonizing asymptomatically the nares of ∼30% of the human population^15^, is no exception.

The intermittent skin colonization by *S. aureus* in healthy individuals (10-20%) clearly contrasts with the nearly persistent colonization of patients with dermo-inflammatory disorders like atopic dermatitis (80-100%)^16^. Interestingly, atopic dermatitis has been associated with several lipid disorders, including defects in sapienic acid, a potent human-specific AFA^17^. It is unclear whether *S. aureus* strains associated with atopic dermatitis are exceptionally impervious to AFAs. The diverse resistance mechanisms used by *S. aureus* against AFAs have been reviewed elsewhere^14^. Notably, the bacterium has long been known to secrete a fatty acid-modifying enzyme (FAME) that mediates AFA-detoxification via esterification with cholesterol or, with lower efficacy, other alcohools^18^. The identity of the protein(s) responsible for FAME activity has remained elusive.

To uncover FAME and other protective strategies against the deleterious effects of AFAs, proteins secreted by *S. aureus* grown in the presence of a subinhibitory concentration of AFAs have been examined^19^. This study revealed that the bacterium boosted its release of the lipolytic lipase Lip2 (also referred to as Geh or Sal2) when primed with AFAs^19^. Recently, we uncovered Lip2 and other lipases as major components of membrane vesicles (MVs) from *S. aureus* irrespective of the presence of AFAs in the growth medium^20^. Given that the impact of *S. aureus* lipases on bacterial susceptibility to AFAs in various lipid environments has never been thoroughly investigated, the protective effects of lipase-loaded MVs against AFAs^20^ prompted us to probe the role of lipases in bacterial adaptation to AFAs.

Here, we unveiled Lip2 as an unanticipated resistance factor against AFAs. Lip2 is necessary and sufficient for the esterification of AFAs to cholesterol, with consequences for bacterial growth in liquid cultures, biofilms, and on mammalian skin.

## Results

### *S. aureus* lipases mediate resistance against AFAs

Our recent proteomics study has uncovered lipases as major components of MVs from *S. aureus* even when the bacterium was grown in the presence of AFAs^20^. These observations suggest that bacteria utilize lipases to cope with AFAs. In agreement with the previously reported protective roles of MVs against AFAs^20^, we hypothesized that lipases are required for bacterial growth in the presence of AFAs.

To test this hypothesis, we monitored the growth kinetics of wild-type USA300 JE2 (WT) or its mutant defective for both Lip1 and Lip2 lipase production (henceforth referred to as Δlip^21^) in a rich medium where Δlip displayed no growth defect (Fig. 1A,B). Notably, even upon treatment with palmitoleic acid (PA), a major AFA of mammalian skin^22^ and nasal fluid^9^, no clear differences in growth behaviors were apparent between Δlip and WT, which were both strongly inhibited by 50 µM PA, i.e., PA concentration in the nasal fluid^9^ (Fig. S1A and Fig. 1A,B). The abundance of PA generally correlates with that of cholesterol in the nasal fluid^9^. Owing to cholesterol-protective roles against AFAs^14^, we wondered whether cholesterol would boost the growth of WT and Δlip in the presence of otherwise inhibitory amounts of PA. Strikingly, cholesterol, which alone does not alter the replication of *S. aureus* (Fig. S1B,C), counteracted PA toxicity in a lipase-dependent manner (Fig. 1A,B). The heightened susceptibility of Δlip to AFAs was readily apparent when a different fatty acid, linoleic acid (LA), was used (Fig. S1D). In experimental settings where WT and Δlip were similarly inhibited by LA, cholesterol was protective only for WT (Fig. S1E). In addition to optical density readings, the lipase-dependent protective effects of cholesterol were also evidenced by CFU (colony forming unit) enumeration (Fig. 1C).

**Figure 1.**
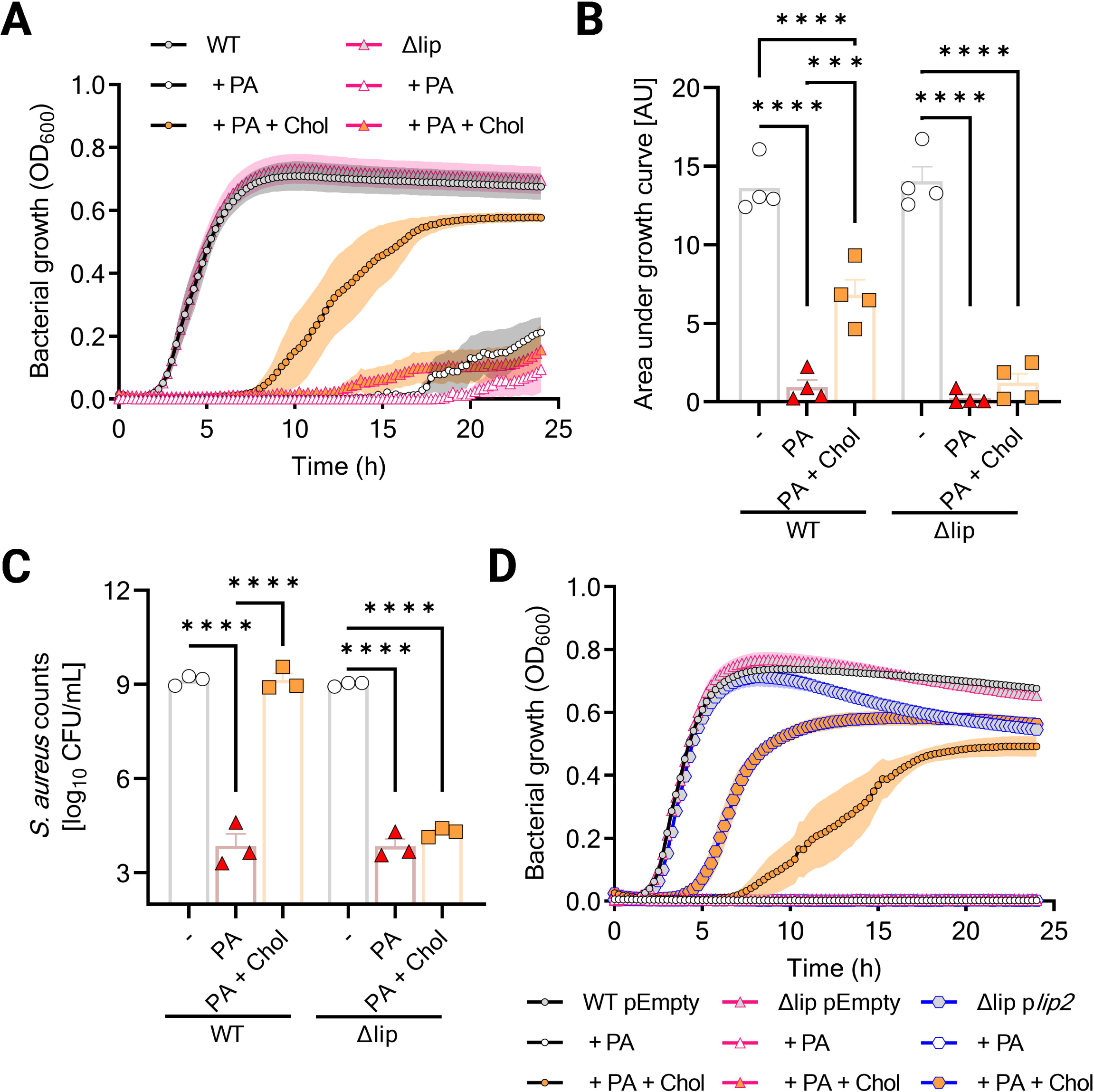
Lipases protect *S. aureus* against palmitoleic acid. **A**, Optical density at 600 nm (OD_600_) was measured over 24 h to monitor the growth of USA300 JE2 (WT) and its Lip1- and Lip2-defective double mutant (Δlip) in plain nutrient broth (NB), or NB supplemented with palmitoleic acid (PA) or PA and cholesterol (Chol). **B**, Area under the growth curves (shown in **A**) was computed in arbitrary units (AU). **C**, Viable WT and Δlip were enumerated upon growth for 24 h in NB, or NB supplemented with PA or PA + cholesterol (Chol). **D**, WT and Δlip bearing an empty plasmid (pEmpty), and Δlip complemented with p*lip2* were grown as described in **C** while OD_600_ was measured. Data are presented as mean ± standard error of the mean (SEM) for 3 (**C**) or 4 (**A**,**B**,**D**) biological replicates. Statistical significance was determined by one-way analysis of variance (ANOVA) with Tukey’s multiple comparisons test. \*\*\**P* = 0.0002, \*\*\*\**P* < 0.0001.

### The lipase Lip2 is sufficient for cholesterol-mediated protection against AFAs

To determine whether Lip1 and Lip2 were both required for the phenotype of the double lipase mutant Δlip or one of both enzymes played a dominant role, we first tested a single *lip2* mutant (Δ*lip2*^23^) and its otherwise isogenic USA300 wild-type strain for growth in the presence of LA. Δ*lip2* displayed a longer lag phase (∼ 11 h) compared to its WT (∼ 7 h), suggesting that Lip2 is protective against LA (Fig. S1E). Next, Δlip was complemented with *lip2* on a plasmid (p*lip2*). The complemented strain Δlip p*lip2* had no growth advantage in rich medium over a Δlip mutant carrying an empty plasmid (pEmpty). However, p*lip2*-complementation enabled Δlip to proliferate in the presence of toxic amounts of PA (Fig. 1D), LA (Fig. S2A), or sapienic acid (SA) (Fig. S2B), albeit only upon addition of cholesterol. The growth defect of Δlip pEmpty in media supplemented with cholesterol and AFAs, as compared to either WT pEmpty or Δlip p*lip2*, was alleviated when this mutant was provided with MVs from WT USA300 (Fig. 2A). MV-associated lipases appeared to be responsible for MV-mediated complementation of Δlip pEmpty since the AFA-resistance was not restored when MVs were from Δlip (Fig. S2C). Importantly, recombinant Lip2 also enabled the growth of Δlip pEmpty upon exposure to AFA and cholesterol (Fig. 2B).

**Figure 2.**
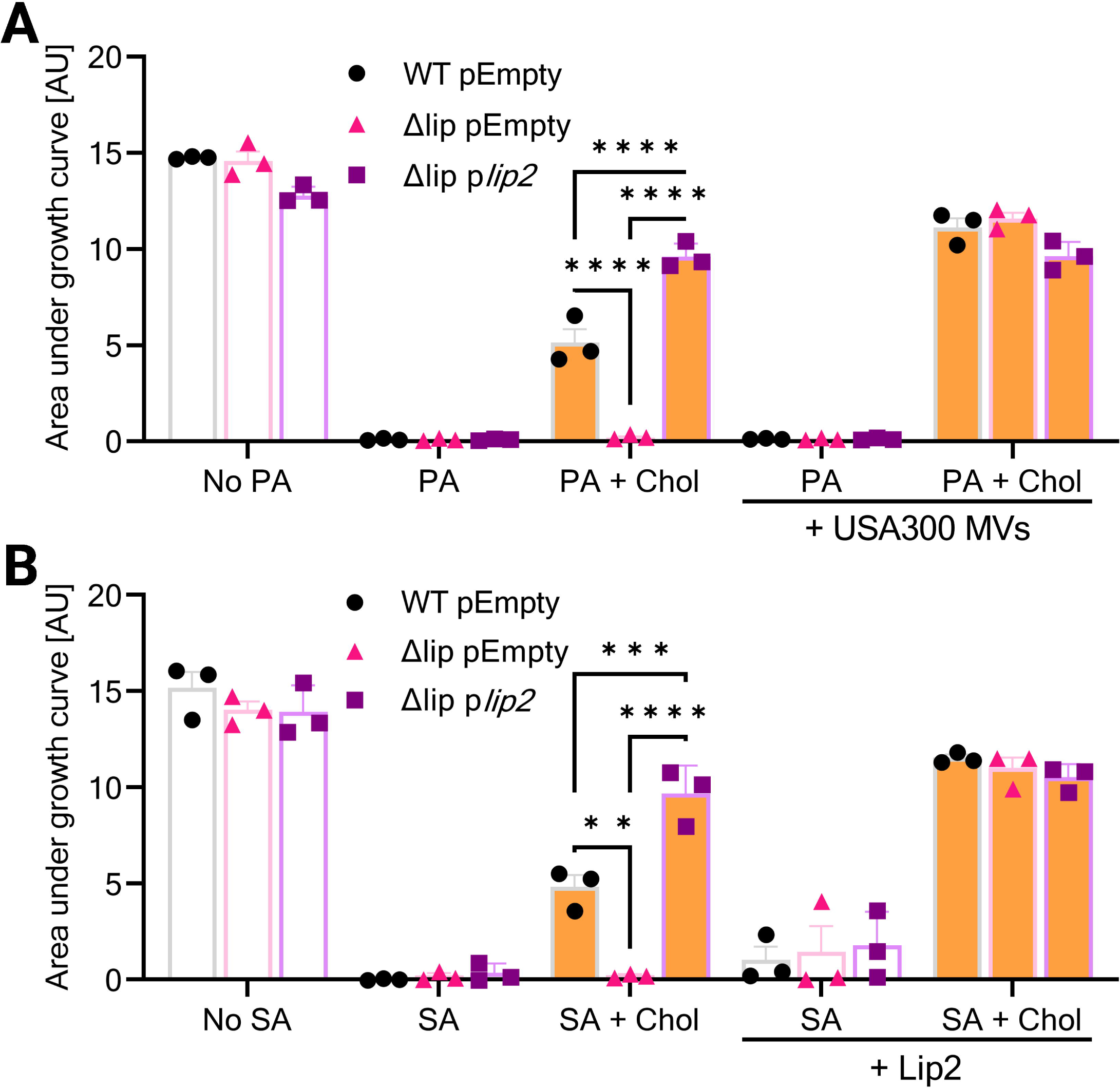
Exogenous lipases enable cholesterol-dependent growth in the presence of AFAs. **A**, Wild-type USA300 JE2 and its isogenic Δlip mutant with pEmpty, and Δlip complemented with p*lip2* were grown in plain NB, or NB with or without USA300 membrane vesicles (MVs) and supplemented with PA or PA + Chol. Computed area under growth curves was plotted. **B**, Area under the curves of the strains described in (**A**) upon growth in NB, or NB supplemented with sapienic acid (SA) or SA + Chol, with or without recombinant Lip2. Data shown are mean + SEM (*n* = 3). Statistical significance was evaluated by two-way ANOVA with Tukey’s multiple comparisons test. \*\**P* = 0.0019, \*\*\**P* = 0.0009, \*\*\*\**P* < 0.0001.

In addition to the prominent role of Lip2 in cholesterol-mediated protection against AFAs, we sought to investigate a possible involvement of Lip1. Therefore, Δlip was complemented with p*lip1*. The generated strain was then tested along with pEmpty-bearing WT and Δlip, as well as p*lip2*-complemented Δlip. In clear contrast to p*lip2*, p*lip1* did not allow Δlip to benefit from cholesterol and thereby grow in the presence of PA (Fig. S3A) or α-linoleic acid (ALA) (Fig. 3A), suggesting that Lip2 is solely responsible for cholesterol-aided protection against AFAs. Next, to test whether the catalytic activity of Lip2 was required to mediate cholesterol-dependent AFA resistance, we genetically engineered p*lip2* into p*lip2*^S412A^, bearing a catalytically inactive copy of Lip2 (Lip2 S412A), as demonstrated in previous studies^13, 23^. Upon complementation with this catalytically inactive form of Lip2, the double mutant Δlip displayed no lipase activity, as assessed with a long-chain fatty acid ester substrate (Fig. S3B). This mutant was also unable to benefit from cholesterol supplementation to grow in the presence of ALA (Fig. 3B) or SA (Fig. S3C). Taken together, our data indicate that Lip2 requires its enzymatic activity to mediate cholesterol-dependent AFA resistance.

**Figure 3.**
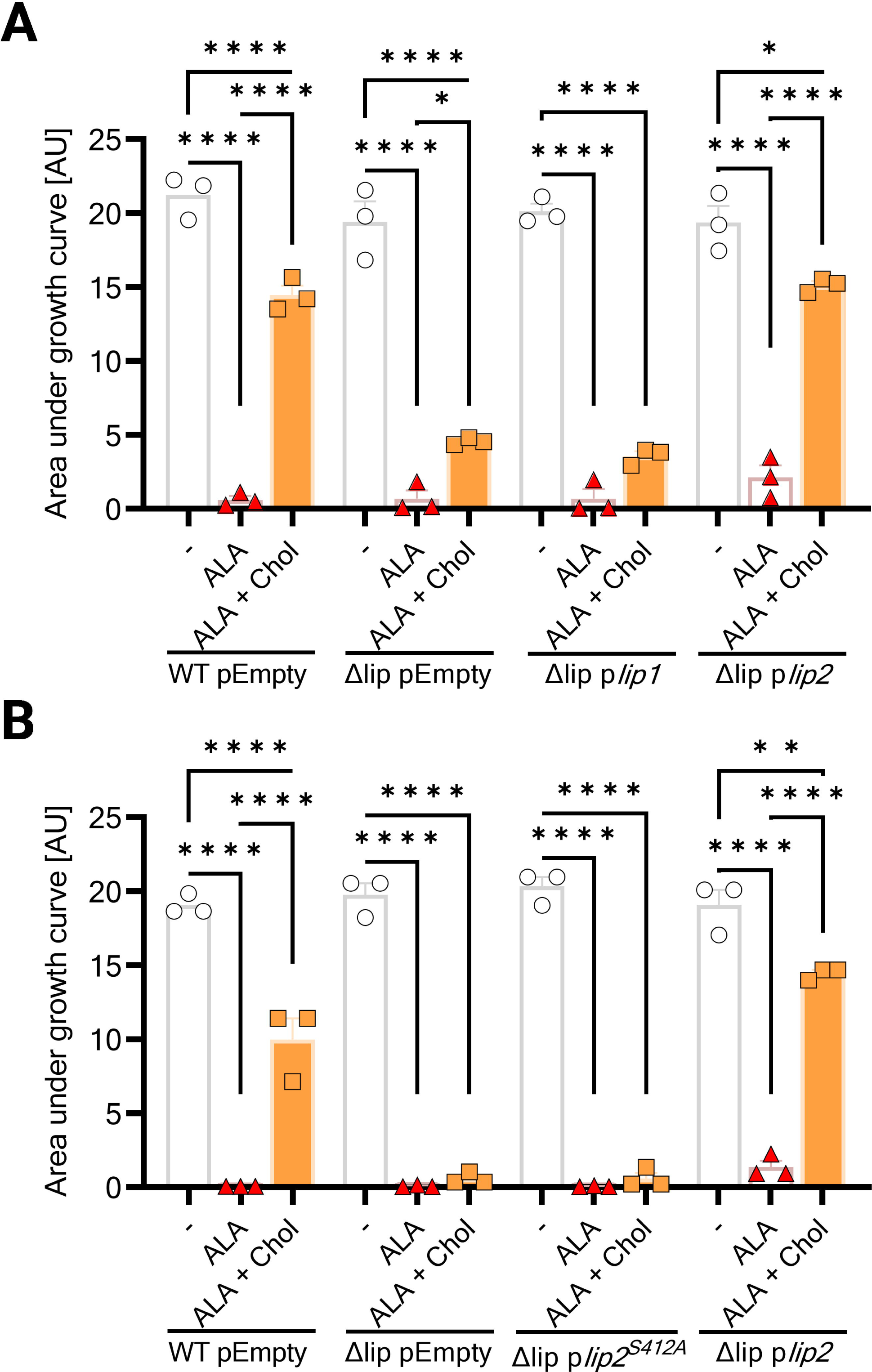
Catalytically active Lip2 is required for cholesterol-mediated resistance to AFAs. **A**, Wild-type USA300 JE2 and its isogenic Δlip mutant with pEmpty, and Δlip complemented with either p*lip1* or p*lip2* were grown for 24 h in basic medium (BM), or BM supplemented with α-linoleic acid (ALA) or ALA + Chol. Computed area under growth curves was plotted. **B**, Area under the curves of pEmpty-bearing wild-type USA300 JE2 and its isogenic Δlip mutant, and Δlip complemented with either p*lip2^S^*^412^*^A^* or p*lip2* cultured for 24 h in BM, or BM supplemented with α-linoleic acid (ALA) or ALA + Chol. Data shown are mean + SEM (*n* = 3). Statistical significance was evaluated by one-way ANOVA with Tukey’s multiple comparisons test. \**P* < 0.05, \*\**P* = 0.0011, \*\*\*\**P* < 0.0001.

### Cholesterol-mediated protection against AFAs is widespread in *S. aureus*

To investigate whether cholesterol was protective against AFAs for *S. aureus* strains other than USA300, USA400 MW2, USA200 UAMS-1, SH1000 and Newman were assessed for growth in the presence of cholesterol and LA. All these *S. aureus* strains clearly benefited from cholesterol to better grow in the presence of LA (Fig. 4A). Interestingly, Newman’s protection by cholesterol (i.e., LA versus LA + cholesterol) failed to reach statistical significance (*P* = 0.1062), in agreement with the fact that Newman Lip2-encoding gene (*lip2*) is disrupted by a prophage^24^. Upon complementation with p*lip2*, Newman became able to replicate in a cholesterol-dependent manner at otherwise toxic LA concentrations (Fig. S4A,B). Surprisingly, p*lip2* imparted a strong metabolic burden to Newman in a rich medium (nutrient broth), which was alleviated by a change of medium. In another rich medium (basic medium), where pEmpty or p*lip2*-bearing Newman grew similarly, the role of Lip2 as a resistance mechanism against AFA became apparent upon growth in planktonic conditions (Fig. 4B and Fig. S4C,D), or within biofilms (Fig. 4C).

**Figure 4.**
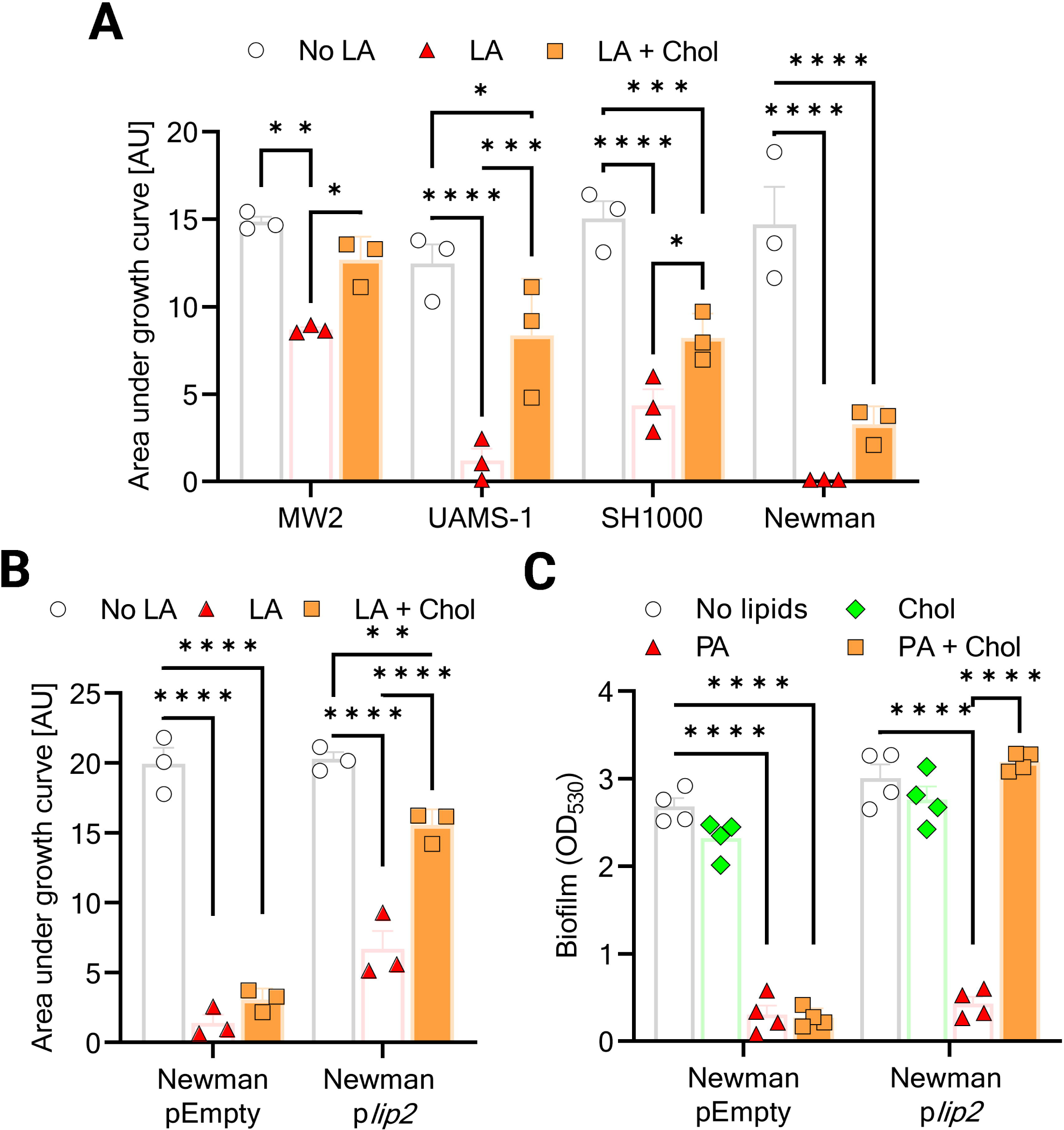
Various *S. aureus* strains utilize cholesterol to resist AFAs. **A**, *S. aureus* strains (MW2, UAMS-1, SH1000, and Newman) were cultured for 24 h in plain NB, or NB supplemented with linoleic acid (LA) or LA and Chol. Growth was computed as area under the curves. **B**, Area under the curves of Newman with either pEmpty or p*lip2* grown for 24 h in plain BM, or BM supplemented with LA or LA + Chol. **C**, Optical density at 530 nm (OD_530_) was measured after safranin staining of biofilms formed for 24 h by Newman with either pEmpty or p*lip2* in plain BM, or BM supplemented with Chol, PA, or Chol + PA. Data shown are mean + SEM for 3 (**A, B**) or 4 (**C**) biological replicates. Statistical significance by two-way ANOVA with Tukey’s multiple comparisons test. \**P* < 0.05, \*\**P* < 0.01, \*\*\**P* < 0.001, \*\*\*\**P* < 0.0001.

### Lipase Lip2 esterifies AFAs

In addition to lipid hydrolysis, lipases catalyse esterification and transesterification^25^. Recently, a secreted lipase of *Vibrio parahaemolyticus* has been shown to esterify cholesterol with host-derived polyunsaturated fatty acids^26^. Moreover, *S. aureus* and many other staphylococci are known to utilize FAME to detoxify unsaturated, antimicrobial fatty acids by esterification with hydroxylated substrates, including cholesterol^18, 27^. The protein responsible for FAME activity has been enigmatic for three decades. In the light of the Lip2-dependent protective effects of cholesterol against AFAs, we tested recombinant Lip2^23^ for FAME (esterification) activity. Upon incubation of Lip2 with LA and cholesterol, followed by lipid extraction and high-performance thin layer chromatography (HPTLC), we detected cholesteryl linoleate, a cholesteryl ester (Fig. 5A). Lip2-catalysed esterification of LA with cholesterol was also confirmed via ultra-high performance liquid chromatography-electrospray ionization-tandem mass spectrometry (UHPLC-MS/MS) (Fig. 5B). Lip2 but not catalytically inactive Lip2 S412A displayed esterifying activity on all five AFAs we tested, irrespective of chain length and degree of unsaturation, as revealed by HPTLC (Fig. S5A).

**Figure 5.**
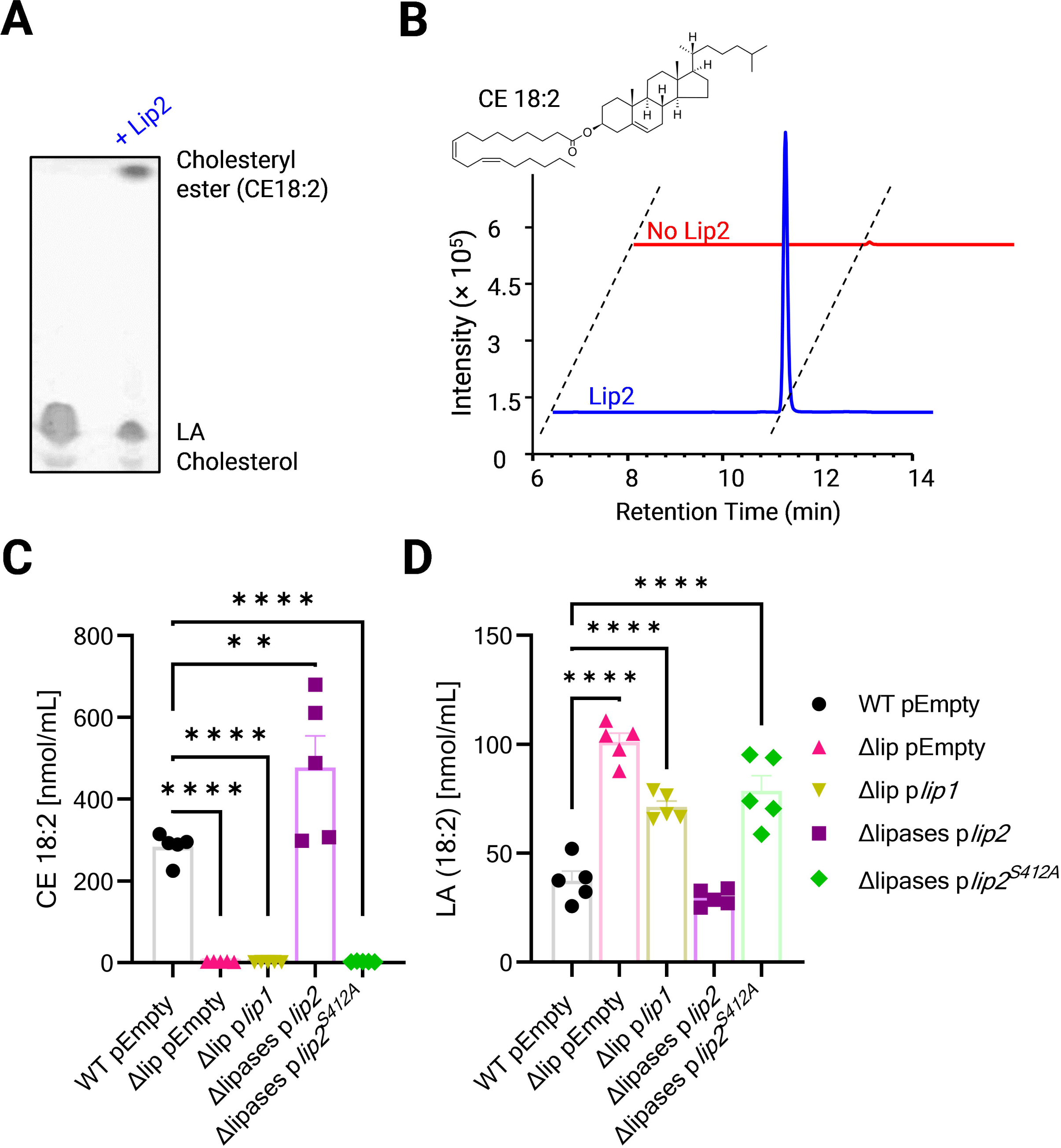
Lip2 lipase mediates esterification of AFAs. **A**, HPTLC of lipid extracts after incubation of LA and Chol with or without recombinant Lip2. **B**, Structure of cholesteryl linoleate (CE 18:2) and representative extracted ion chromatograms of *m/z* 648.585±0.010 (precursor type [M+NH4]^+^ in positive ion mode of CE 18:2) demonstrating detection of CE 18:2 upon co-incubation of LA, Chol and recombinant Lip2. **C-D**, UHPLC-MS/MS lipid analysis upon incubation of *S. aureus*-conditioned media from the indicated strain (WT pEmpty, Δlip pEmpty, Δlip p*lip1*, Δlip p*lip2*, or Δlip p*lip2^S412A^*) with Chol and LA. CE 18:2 (**C**) and LA (**D**) were measured. Bar graphs (**C**, **D**) are means + SEM for five biological replicates. Statistical significance by one-way ANOVA with Dunnett’s test relative to WT pEmpty. \*\**P* = 0.0034, \*\*\*\**P* < 0.0001.

Further, to demonstrate that Lip2 released by *S. aureus* can esterify cholesterol with AFAs, we treated *S. aureus*-conditioned media from plasmid-bearing Δlip or WT with AFAs and cholesterol prior to lipid analysis. Cholesteryl esters (CE) were detected only in Lip2-expressing WT pEmpty and Δlip *plip2* strains by HPTLC (Fig. S5B) or UHPLC-MS/MS (Fig. 5C). Accordingly, CE production was concomitant with decreased concentrations of free AFA (Fig. 5D) and cholesterol (Fig. S5C). Taken together, our data identify Lip2 as FAME, which detoxifies AFAs by esterification with cholesterol.

Despite a clear preference for cholesterol, FAME has also been shown to use other alcohols for AFA esterification^18^. Consistent with Lip2-mediated FAME activity, CE were still produced by Lip2-expressing strains, when *S. aureus*-conditioned media were supplemented with approximately eight hundred times molar excess of ethanol to compete with cholesterol for AFA esterification (Fig. S5D). Moreover, this experimental setup unveiled that while Δlip p*lip2* or WT pEmpty esterified AFA with either ethanol or cholesterol, Δlip complemented with p*lip1* esterified AFA with ethanol only (Fig S5D), suggesting FAME activity for Lip1 with ethanol and presumably other alcohols as substrates. The requirement of Lip2 for cholesterol esterification was also evidenced in the USA400 strain, MW2 WT, in which single mutants defective in Lip1 (MW2 Δ*lip1*) or Lip2 (MW2 Δ*lip2*) were generated. Conditioned medium by MW2 Δ*lip2* could esterify ethanol (Fig. S5F) but not cholesterol (Fig. S5E,F), while MW2 Δ*lip1*-conditioned medium retained the MW2 WT’s ability to utilize ethanol and cholesterol for AFA esterification (Fig. S5E,F). Together, these data underline cholesterol as preferred substrate for Lip2-mediated esterification of AFAs. As exemplified by Lip1, the ability to modify AFAs with alcohols is a poor predictor for cholesterol utilization and likely explains why FAME has remained elusive for so long.

### Membrane damages caused by AFAs are not prevented by cholesterol

Our data (Fig. 5 and Fig. S5) strongly suggest that the protective effects of cholesterol against AFAs are due to the Lip2-mediated esterification/detoxification of AFAs with cholesterol. However, additional or alternative mechanisms might contribute to our observations. For instance, Lip2-dependent binding to cholesterol could lead to the formation of bacterial aggregates with decreased susceptibility to AFAs. To assess this possibility, we used dehydroergosterol (DHE) as a fluorescent cholesterol analogue^28^ for binding assays with pEmpty-bearing USA300 WT and Δlip, as well as Δlip complemented with p*lip1*, p*lip2* or p*lip2^S^*^412^*^A^*. We did not observe any difference between Lip2-defective and Lip2-proficient strains in their ability to bind sterols (Fig. S6). This suggests that impaired cholesterol binding is unlikely to be the reason why Lip2-deficient *S. aureus* failed to utilize cholesterol against AFAs.

Despite similar binding to sterols, it remained plausible that Lip2-deficient bacteria were defective in preventing interactions with AFAs in the presence of cholesterol. We took advantage of a palmitoleic acid analogue (PA alkyne) and click chemistry with azide fluor 488 for AFA-binding studies^20, 29^ with or without cholesterol supplementation. We found that, irrespective of Lip2 expression and despite cholesterol treatment, PA alkyne clearly bound to WT and mutants, as revealed by fluorometry (Fig. S6B) and flow cytometry (Fig. S6C). Thus, our results suggest that cholesterol does not prevent *S. aureus* membrane-targeting by AFAs.

Another putative protective mechanism of cholesterol could be to preserve the membrane integrity of Lip2-expressing bacteria in the presence of AFAs, which would be reminiscent of the role of the golden carotenoid pigments staphyloxanthin in *S. aureus*^30^. Since membrane-damaging effects of AFAs include loss of membrane potential^31^, we examined the membrane potential of WT and mutants upon treatment with PA, or PA and cholesterol. PA-treated bacteria displayed an almost undetectable membrane potential, which was not restored by co-treatment with cholesterol (Fig. S6D). This suggests that cholesterol per se does not prevent membrane damages caused by AFAs. Thus, Lip2-mediated esterification of AFAs which cholesterol seems to be the only mechanistical explanation of the protective effects of cholesterol towards AFAs.

### Lip2 is a conserved protein that can be disrupted by prophages

To gain unprecedented insights into a potential involvement of Lip2 into tissue tropism, we delved into our custom database of almost four thousand genomes of *S. aureus* obtained from the Bacterial and Viral Bioinformatics Resource Center (BV-BRC)^32^ to identify potential association of the presence or absence of intact lip2 with specific *S. aureus* clones or specific human habitats. This database encompasses blood (1481), nose (1587), and skin (767) isolates. An *in silico* polymerase chain reaction^33^ was used to retrieve sequences of *lip2* in 91.23% (1352 out of 1481), 88.78% (1409 out of 1587), or 95.2% (730 out of 767) of blood, nose, or skin isolates, respectively (Fig. 6A). Next, Lip2 protein sequences were deduced from *lip2* genes. In keeping with the widespread presence of lipases in staphylococci^34^, Lip2 appeared to be highly conserved in *S. aureus* strains irrespective of the isolation site (Fig. S7). Interestingly, accross the seven major sequence types (ST) of our database, the ST dictated Lip2 diversity (Fig. S8). Irrespective of ST, eight mutation hotspots were apparent in Lip2 (Fig. S9A), with some mutations cooccurring in several clonal groups (Table S1). It remains to be elucidated whether these modifications impact Lip2 lipase/FAME activity.

**Figure 6.**
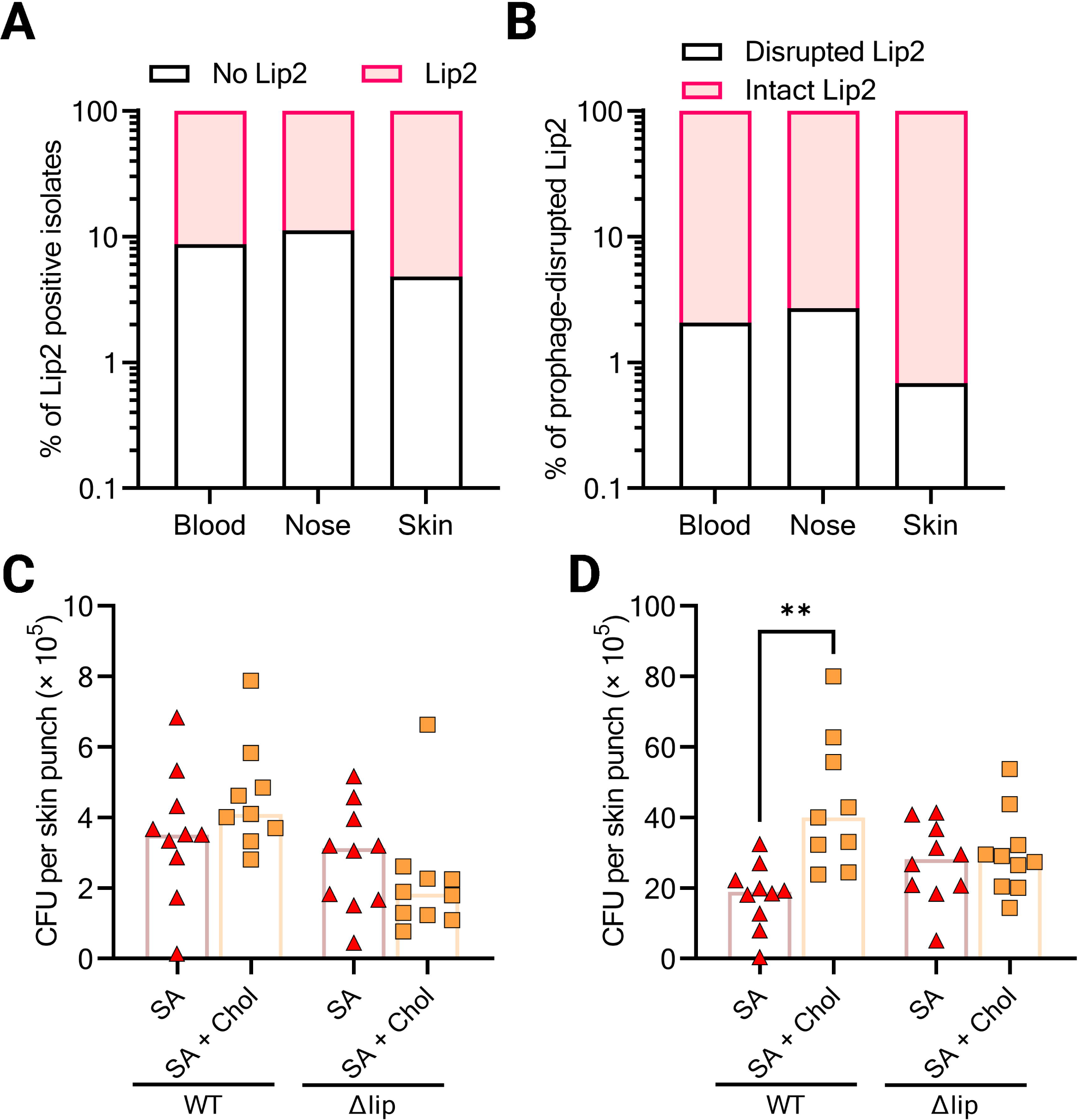
The capacity to manipulate cholesterol governs skin colonization by S. aureus. **A**, The occurrence of *lip2*, as detected via *in silico* PCR, is displayed according to the isolations site for *S. aureus* genomes in our database. **B**, The sequences of *lip2* (retrieved in **A**) were analysed for prophage bearing. **C**-**D**, USA300 JE2 (WT) and its isogenic Δlip mutant were used to topically colonize the skin of mice co-treated with cholesterol (Chol) and/or sapienic acid (SA). Five mice per group correspond to 9 or 10 skin punches, which were strongly vortexed to dislodge surface-attached bacteria (**C**), and then minced to release bacteria located in the deeper skin tissue (**D**). Viable bacteria were counted as colony forming units (CFU). Bar graphs (C and D) are medians. Statistical significance was evaluated by Kruskal-Wallis test with Dunn’s multiple comparisons. \*\**P* = 0.0013.

The nucleotide sequence of *lip2* encompasses a conserved integration site for prophages. A disruption of *lip2* gene by a prophage inactivates Lip2^13^. Therefore, we had a second look at *lip2* sequences to investigate how often prophage insertion occurred. Strikingly, only 2% (71 out of 3491) of the strains exhibited a prophage-disrupted *lip2*. Roughly half of the strains with prophage-disrupted *lip2* were from the sequence type ST398, a livestock-associated *S. aureus* lineage, which represents only 8% of the genomes in our database (Fig. S9B,C). Remarkably, prophage-mediated inactivation of *lip2* was more frequent in blood and nose isolates (2.1% and 2.7%, respectively) than in skin isolates (0.7%) (Fig. 6B). These results suggested that an intact *lip2* may be required for successful skin colonization.

### Skin colonization by *S. aureus* is governed by environmental lipids

To ascertain the requirement of lipases for skin colonization *in vivo*, we opted for a well-established mouse skin colonization model^21, 35, 36^, which mimics human atopic dermatitis. This model leverages the impaired skin barrier function upon extensive tape-stripping to improve skin colonisation by *S. aureus* in a similar manner as in human atopic dermatitis patients. The tape-stripped skin was topically colonized with *S. aureus*. With such a model, we previously observed that wild-type USA300 JE2 and Δlip did not differ in their capacity to colonize mouse skin^37^. Since tape-stripping is known to deplete lipids from the skin^38, 39^, we repleted mouse skin with sapienic acid (SA), or cholesterol plus SA during colonization with either WT or Δlip. Whereas skin colonization by Δlip was largely unaffected by cholesterol application, Lip2-proficient WT appeared to benefit from cholesterol to better colonize the skin in the presence of SA (Fig. 6C,D). Taken together, our results strongly suggest that *S. aureus* utilizes its lipases to manipulate environmental lipids and proliferate on the skin.

## Discussion

The immense success of *S. aureus* as an opportunistic pathogen requires strategies to circumvent host defences, including AFAs^8, 40^. The huge variety of the resistance mechanisms used by bacteria against AFAs strongly suggests a key role for AFAs at the host – pathogen interface^14^. Importantly, bacteria utilize a vast array of lipases to hydrolyse lipids in their environment with sometimes fatal consequences for microbial competitors^12, 41^ or eukaryotic host cells^42^. Bacteria-mediated lipid hydrolysis releases long-chain fatty acids, which can be toxic to microbes^12, 23, 43^. For *S. aureus* and other staphylococci, it is currently thought that lipase-expressing strains utilise FAME to detoxify AFAs released by lipases. However, the identity of the protein(s) responsible for FAME activity has remained elusive for over three decades. Here, we uncovered that lipases are responsible of FAME activities in *S. aureus*. While both lipases Lip1 and Lip2 use ethanol and likely other alcohols for AFA esterification, only Lip2 esterifies AFAs with cholesterol. The ability to utilise cholesterol proved vital as cholesterol protected Lip2-proficient strains against AFA toxicity in planktonic as well as biofilm settings. The unanticipated substrate flexibility of Lip2 strongly suggests a more complex role for bacterial lipases in shaping the host lipid landscape than previously thought, with potential consequences for the microbiome.

The production of lipases by *S. aureus* was first documented more than a century ago^44^. Ever since, evidence of the requirement for bacterial lipases during *S. aureus* infection has been accumulating. For instance, anti-lipase IgG antibodies have been detected in patients infected with *S. aureus*^45^. Furthermore, the expression of lipase-encoding genes has been demonstrated during *S. aureus* infection in a murine renal abscess model^46^. However, only a handful of studies could show diminished virulence for lipase-deficient mutants in mice infected with *S. aureus*^13, 34, 47^. Moreover, numerous studies have used strains with prophage-disrupted-*lip2* to successfully establish murine models of infection with *S. aureus*^8, 48^. In sum, while it is reasonable to perceive lipases as virulence factors, rigorous testing in various models is still needed to fully understand the role played by *S. aureus* lipases during colonization/infection. Our data suggests that suitable environmental lipids are needed to illuminate the versatility of *S. aureus* lipases.

In *S. aureus*, the expression of lipase-encoding genes is controlled by the global regulators Agr and SarA^49, 50^. Accordingly, the secretion of lipases Lip1 and Lip2 is impaired in mutants defective for Agr and/or SarA^51^. In a similar manner, FAME production is drastically impaired in mutants deficient in Agr or SarA^52^. In addition to a rather similar regulation, a strong correlation between lipase and FAME activities, i.e., esterification of fatty acids, has been observed for *S. aureus*^18^ and some coagulase-negative staphylococci^27^. Moreover, Kumar and co-workers uncovered that media conditioned by *S. aureus* strains with high lipolytic activity led to profound changes in the bovine heart lipids, including the production of cholesteryl esters^53^. Our study provides evidence that Lip2 is the lipase catalysing the esterification of AFAs with cholesterol. Lip2 can also use ethanol for AFA esterification whereas Lip1-mediated esterification of AFAs only took place with ethanol. This mirrors the substrate preference of Lip1 and Lip2 for short-chain and long-chain fatty acids, respectively^23, 44^. It is yet unclear which structural features dictate substrate preference and activity in *S. aureus* lipases. We surmise that these features also govern the utilisation of cholesterol by Lip2, which could represent a novel therapeutic target.

Collectively, with our newfound understanding of Lip2 activities, it is enticing to posit that staphylococcal lipases play an underappreciated role in shaping host-derived lipids on the skin and at mucosal surfaces. Eavesdropping on the lipid-mediated crosstalk between microbiomes and hosts could prove pivotal for a better understanding and prevention of colonization by opportunistic pathogens.

## Methods

### Bacterial strains and growth conditions

Bacterial strains and plasmids used in this study are detailed in Table S2. *S. aureus* and *Escherichia coli* strains were routinely grown overnight at 37°C in tryptic soy broth (TSB) or lysogeny broth (LB), respectively. Whenever appropriate, the medium was supplemented with ampicillin (100 µg/mL), kanamycin (30 µg/mL), or chloramphenicol (10 µg/mL).

### Construction of strains

Primers used are listed in Table S3. In-frame deletion of *lip1* or *lip2* was performed with pIMAY as described previously^54^. Gene deletion was confirmed by PCR and sequencing. For mutant complementation experiments, empty pALC2073^55^ (pEmpty), pALC2073-*lip1* (p*lip1*), and pALC2073-*lip2*^23^ (p*lip2*) were used. To generate p*lip2^S^*^412^*^A^*, p*lip2* was amplified with mutagenic primers. *E. coli* IM08 was then transformed with the DpnI-treated PCR product. After plasmid purification, successful mutagenesis was confirmed by digestion with PaeI and sequencing.

### Purification of recombinant lipases

N-terminally His_6_-tagged Lip2 or Lip2 S412A^23^ was overexpressed in *E. coli* BL21 (DE3). After cell lysis, recombinant protein was purified using nickel resin according to standard procedures^13^.

### Membrane vesicle purification

MVs were isolated with the ExoQuickTC reagent (EQPL10TC; System Bioscience) as described elsewhere^20, 56^. Briefly, bacteria grown overnight were diluted to an optical density at 600 nm of 0.1 (OD_600_) in 20 ml fresh TSB and grown with shaking for 6 h (late exponential growth phase). After centrifugation, supernatants were sterile filtered and concentrated with 100-kDa centrifugal concentrator cartridges (Vivaspin 20; Sartorius) prior to precipitation with ExoQuickTC and resuspension in phosphate-buffered saline.

### Growth assays

Growth assays were performed in TSB (Oxoid), nutrient broth no.2 (NB; Oxoid) or basic medium (BM: 1% soy peptone, 0.5% yeast extract, 0.5% NaCl, 0.1% glucose and 0.1% K_2_HPO_4_, pH 7.2) as described previously^20^. Overnight bacterial cultures were diluted to an OD_600_ of ∼0.01 in plain medium or medium supplemented with AFAs (50 to 200 µM), cholesterol (50 to 100 µM), MVs (1 µg/mL), and/or recombinant Lip2 (1 µg/mL). Bacteria were then grown in a 96-well plate (U-bottom) at 37°C with linear shaking at 567 cpm (3-mm excursion) for 24 h. The OD_600_ was measured every 15 min with an Epoch 2 plate reader (BioTek). Areas under growth curves were computed with GraphPad Prism 9.5.1.

### Biofilm assay

Biofilms formed under static conditions at 37°C for 24 h in cell culture 24-well plates (Greiner) were stained with safranin as described elsewhere^57^. Unbound safranin was washed with PBS, and biofilm-associated safranin was incubated with 70% ethanol and 10% isopropanol for elution. A CLARIOStar microplate reader (BMG Labtech) was used to measure OD_530_ and quantify biofilms.

### Lipase activity assay

The lipase activity of bacteria-conditioned media was assayed with *para*-nitrophenyl palmitate as previously described^23^. Bacteria-conditioned media were diluted fifty times with the assay buffer (50 mM Tris·HCl, 0.005% Triton X-100, 1 mg/mL gum arabic at pH 8.0) supplemented with 0.8 mM *para*-nitrophenyl palmitate. After incubation at 37°C for 30 minutes, OD_405_ was measured with a CLARIOStar microplate reader (BMG Labtech).

### FAME activity assay, lipid extraction and HPTLC

Recombinant lipases or bacteria-conditioned media were diluted in 0.1 M sodium phosphate buffer (pH 6) supplemented with AFAs and cholesterol. Upon overnight incubation in glass vials at 37°C with shaking, methanol (MeOH) and chloroform were added to stop the reaction and extract lipids according to the Bligh and Dyer protocol^58^. The organic fraction was transferred to a fresh vial, dried, and resuspended in 2:1 (vol/vol) chloroform: MeOH. Lipid extracts were then applied to silica gel high-performance thin-layer chromatography (HPTLC) plates (silica gel 60 F_254_, Merck) using a Linomat 5 sample application unit (CAMAG). Plates were developed in an automatic developing chamber ADC 2 (CAMAG) with a mobile phase system 90:10:1 (vol/vol/vol) petroleum ether: ethyl ether: acetic acid^26^. Lipid spots were visualized in an iodine vapor chamber.

### Internal standards and chemicals used for lipid analysis by untargeted UHPLC MS/MS

EquiSPLASH™ LIPIDOMIX® quantitative mass spectrometry internal standard, phosphatidic acid 15:0-18:1 (d7), cholesterol (d7), cholesteryl ester (CE) 18:1 (d7), lyso sphingomyelin (LSM) d18:1 (d9) and palmitoyl-L-Carnitine (CAR) 16:0 (d3) were obtained from Avanti Polar Lipids (Alabaster, AL, USA). Arachidonic acid (AA) (d11) and ceramide (Cer) d18:1-15:0 (d7) were purchased from Cayman Chemicals (Ann Arbor, MI, USA). Isopropanol (IPA), acetonitrile (ACN) and methanol (MeOH) in Ultra LC-MS grade were from Carl Roth (Karlsruhe, Germany). Ammonium formate, formic acid and IPA in HPLC grade were purchased from Merck (Darmstadt, Germany). Purified water was produced by Elga Purelab Ultra (Celle, Germany).

### Sample preparation for lipid analysis by UHPLC MS/MS

Prior to lipid extraction, a mixture of internal standards was prepared by mixing ice-cold MeOH with LIPIDOMIX®, phosphatidic acid 15:0-18:1 (d7), cholesterol (d7), CE 18:1 (d7), LSM d18:1 (d9), CAR 16:0 (d3), AA (d11), and Cer d18:1-15:0 (d7). This internal standard mixture (225 µL) was then added to each sample. Lipid extraction was then performed according to a biphasic extraction method^59, 60^. Samples supplemented with standards were vortexed for 10 s. Next, 750 µL ice-cold methyl tert-butyl ether (MTBE) was added to each sample. After 1h-incubation on ice, each sample was supplemented with water (185 µL) to obtain a final ratio of 10:3:2.5 (vol/vol/vol) for MTBE, MeOH, and water, respectively. Samples were then incubated at room temperature for 10 min to induce phase separation. The upper (organic) phase was transferred to a fresh tube. MTBE:MeOH:water (10:3:2.5; vol/vol/vol) was added to the lower (water) phase for re-extraction of lipids. The upper phase from the second extraction was then combined with the upper phase from the first extraction. The combined extracts were evaporated to dryness with GeneVac EZ2 evaporator (Ipswich, UK) under nitrogen protection. Lipid films were reconstituted in 100 μL MeOH. After vortexing (10 s), sonication (2 min), and centrifugation (10 min, 3,500 × *g*), lipid extracts were transferred to autosampler vials.

A pooled quality control (QC) sample was prepared by mixing 15 μL of each re-constituted sample.

### Lipid analysis by UHPLC MS/MS

Samples were analysed with an Agilent 1290 Infinity UHPLC system (Agilent, Waldbronn, Germany) equipped with a binary pump, a PAL-HTX xt DLW autosampler (CTC Analytics AG, Switzerland) and coupled to a SCIEX TripleTOF 5600 + quadruple time of flight (QTOF) mass spectrometer with a DuoSpray Source (SCIEX, Ontario, Canada). The chromatographic separation was performed on an ACQUITY UPLC CSH C18 column (100 mm × 2.1 mm; particles: 1.7 μm; Waters Corporation, Millford, MA, USA) with precolumn (5 mm × 2.1 mm; 1.7 μm particles). The column temperature was 65°C with a flow rate of 0.6 mL/min. Mobile phase A was composed of water: acetonitrile (2:3; vol/vol) supplemented with 10 mM ammonium formate and 0.1% formic acid (vol/vol). The mobile phase B was IPA:ACN:water 90:9:1 (vol/vol/vol) containing 10 mM ammonium formate and 0.1% formic acid (vol/vol). A gradient elution started from 15% B to 30% B in 2 min, followed by increase of B to 48% in 0.5 min. Mobile phase B was then further increased to 82% at 11 min and quickly reached 99% in the next 0.5 min, followed by holding this percentage for another 0.5 min. Afterwards, the percentage of B was switched back to starting conditions (15% B) in 0.1 min to re-equilibrate the column for the next injection (2.9 min).

UHPLC-MS/MS experiments were operated in both positive and negative mode with injection volumes of 3 µL for positive and 5 µL for negative mode. An MS full scan experiment with mass range *m/z* 50 to 1,250 was selected, while different SWATH windows were acquired for MS/MS experiments (Table S4). The ion source temperature was set to 350°C with curtain gas, nebulizer gas and heater gas pressures 35 lb/in^2^, 60 lb/in^2^, and 60 lb/in^2^, respectively, for both modes. The ion spray voltage was set to 5,500 V in the positive mode and −4,500 V in negative mode. The declustering potential was adjusted to 80 V and −80 V for positive and negative polarity mode, respectively. The cycle time was always 720 ms. The collision energy and collision energy spread for each experiment are specified in detail Table S4.

The sequence was started with three injections of internal standard mixture as system suitability test followed by blank extract and QC sample. The whole sequence was controlled by injection of QC sample after every five samples to monitor the performance of the instrument throughout the analytical batch.

### Genomic analyses

For our custom database, *S. aureus* genomes (3,835) downloaded from the BV-BRC^32^. After manually curating the metadata, the database was stratified to blood (1,481), nose (1,587) and skin (767) according to isolation sites. To extract *lip2* gene sequence, *in-silico* PCR was performed with a Perl script (https://github.com/egonozer/in_silico_pcr) using forward and reverse primers 5’-ATGTTAAGAGGACAAGAAGAAA-3’ and 5’-TTAACTTGCTTTCAATTGTGTT-3’, respectively, and allowing 5 mismatch/indels. To detect prophages, results of the *in-silico* PCR were uploaded as one multiFASTA file to PHASTER (https://phaster.ca/)^61^. Prophage-disrupted *lip2* amplicons were not included in the following analysis pertaining to amino acid sequence variability of Lip2. Extracted sequences from the *in-silico* PCR were translated into full-length Lip2 protein sequences, an aligned using MAFFT (v7.310)^62^ with default parameters using Lip2 sequence from *S. aureus* USA300 strain FPR3757 (accession number NC_007793.1) as a reference. A Python script (https://github.com/AhmedElsherbini/Align2XL) was then used to extract mutation rates from the aligned protein sequences.

### Mouse experiments

C57BL/6 mice were colonized epicutaneously with *S. aureus* following tape-stripping as described previously^21, 35, 36^. Briefly, overnight cultures of USA300 JE2 or its isogenic Δlip mutant were washed twice with PBS and adjusted to 5 × 10^9^ cells per mL. An inoculum of 15 µL from the washed bacterial suspension was added to a film paper disc. In addition to bacteria, these discs were supplemented with cholesterol (7 µg) and/or sapienic acid (5 µg). Two discs with bacteria and lipids per mouse were placed onto the back skin that had been shaved and tape-stripped seven times to facilitate *S. aureus* establishment. Finn chambers on Scanpor (Smart Practise, Phoenix, AZ, USA) and plasters (Tegaderm) were used to fix discs on mouse back skin. After 24 hours with frequent monitoring, Finn chambers were removed, mice were euthanized, and a biopsy puncher was used to collect *S. aureus*-colonized skin. These skin punches were vortexed in PBS for 30 s to dislodge surface-attached bacteria. Skin punches were then minced with scalpels and homogenized by vortexing for 30 s in PBS to release tissue-associated bacteria. Surface associated and tissue-associated bacteria were enumerated following serial dilution with PBS, plating on tryptic soy agar, and incubation overnight at 37°C.

### Statistical analysis

Statistical tests, which are all specified in the figure legends, were performed with Prism 9.5.1 (GraphPad), and *P* values < 0.05 were considered significant. Analysis of variance (ANOVA) with Dunn’s, Dunnett’s, Šídák’s, or Tukey’s multiple-comparison test was used.

### Ethics statement

All experimental procedures involving mice were carried out according to protocols approved by the Animal Ethics Committees of the Regierungspräsidium Tübingen (IMIT3/18).

## Acknowledgements

We thank David E. Heinrichs (University of Western Ontario), Paul Fey (University of Nebraska Medical Center), and Friedrich Götz (University of Tübingen) for providing us with bacterial strains. We are indebted to Dr Libera Lo Presti (University of Tübingen) for critical feedback on the manuscript, and to Ulrike Redel for technical support. We acknowledge support by the High Performance and Cloud Computing Group at the Zentrum für Datenverarbeitung (University of Tübingen), the state of Baden-Württemberg through bwHPC and the Deutsche Forschungsgemeinschaft (DFG) through the grant INST 37/935-1 FUGG.

A.K.T. is recipient of a fellowship from the Alexander von Humboldt Foundation. X.F. gratefully acknowledges the support from the China Scholarship Council (grant number 201908080155). This work was supported by grants from the DFG via the Cluster of Excellence EXC 2124 ‘Controlling Microbes to Fight Infections’ project ID 390838134 to A.K.T., B.S, and A.P.

---

**Figure S1. Cholesterol-dependent protective roles of S. aureus lipases against AFAs.**

**A**, USA300 JE2 (WT) and its Lip1- and Lip2-defective double mutant (Δlip) were grown in nutrient broth (NB) supplemented with 0 to 50 µM palmitoleic acid (PA). Computed area under growth curves was plotted. **B-C**, Area under the curves of WT and Δlip upon growth in NB (**B**) or tryptic soy broth (TSB) (**C**) supplemented with 0, 50, or 100 µM cholesterol (Chol). **D**, Area under the growth curves of WT and Δlip in NB or NB plus linoleic acid (LA). **E**, Optical density at 600 nm (OD_600_) was measured over 24 h to monitor the growth of WT and Δlip in TSB, or TSB supplemented with 200 µM LA or 200 µM LA and 100 µM cholesterol (Chol). **F**, The growth of the Lip2 mutant (Δ*lip2*) or isogenic wild-type USA300 (WT) was monitored over 24 h by OD_600_ readings in NB or NB supplemented with 200 µM LA. Data shown are mean ± SEM for at least three biological replicates. Statistical significance was evaluated by two-way ANOVA with Šídák’s multiple comparisons test. \*\**P* = 0.0049.

**Figure S2. The lipase Lip2 is required for cholesterol-mediated protection against AFAs.**

**A,** Wild-type USA300 JE2 (WT) and its isogenic Δlip mutant bearing pEmpty, and Δlip complemented with p*lip2* were grown in plain NB, or NB supplemented with 150 µM LA or 150 µM LA and 75 µM Chol. Growth was computed as area under the curves. **B-C,** Area under the curves of the strains described in A upon growth in NB, or NB supplemented with 50 µM sapienic acid (SA), 50 µM SA + 50 µM Chol (B), or 50 µM palmitoleic acid (PA) + 50 µM Chol and in the presence of membrane vesicles (MVs) from WT or Δlip (C). Data shown are mean + SEM for three **(A)**, four **(B)** or five **(C)** biological replicates. Statistical significance was evaluated by one-**(A, B)** or two-way **(C)** ANOVA with Tukey’s multiple comparisons test. \**P* < 0.05, \*\**P* < 0.01, \*\*\**P* < 0.0006, \*\*\*\**P* < 0.0001.

**Figure S3. Inactivation of Lip2 abrogates cholesterol protection against AFAs.**

**A**, Wild-type USA300 JE2 and its isogenic Δlip mutant with pEmpty, and Δlip complemented with either p*lip1* or p*lip2* were grown for 24 h in plain BM, or BM supplemented with 100 µM PA or 100 µM PA + 100 µM Chol. Computed area under growth curves was plotted. **B**, The *S. aureus*-conditioned media from the indicated strain (WT pEmpty, Δlip pEmpty, Δlip p*lip1*, Δlip p*lip2*, or Δlip p*lip2^S^*^412^*^A^*) were incubated with *para*-nitrophenyl palmitate (pNP-16:0). The release of *para*-nitrophenol, indicative of lipase activity, was quantified by measuring OD_405_. **C**, Area under the curves of pEmpty-bearing wild-type USA300 JE2 and its isogenic Δlip mutant, and Δlip complemented with either p*lip2^S^*^412^*^A^* or p*lip2* cultured for 24 h in BM, or BM supplemented with 75 µM SA or 75 µM SA + 75 µM Chol. Shown are mean + SEM for at least three biological replicates. One-way ANOVA with Tukey’s multiple comparisons test (**A**, **C**) or Dunnett’s test relative to WT pEmpty was used to calculate statistical significance (**B**). \*\*\*\**P* < 0.0001.

**Figure S4. Complementation of Lip2-defective Newman strain by USA300 Lip2.**

**A**, OD_600_ was measured over 24 h to monitor the growth of *S. aureus* Newman with either pEmpty or p*lip2* in NB, or NB supplemented with 100 µM LA or 100 µM LA + 100 µM Chol. **B**, Computed area under the growth curves shown in **A**. **C**, The growth of Newman pEmpty or p*lip2* was monitored over 24 h by OD_600_ readings in basic medium (BM), or BM supplemented with 50 µM PA or 50 µM PA + 50 µM Chol. **D**, Growth curves shown in **C** were computed as area under the curves. Data represented are means ± SEM; *n* = 4 (**A**, **B**) or 3 (**C**, **D**). Statistical significance by one-way ANOVA with Tukey’s multiple comparisons test. \**P* = 0. 0418, \*\**P* = 0.0027, \*\*\*\**P* < 0.0001.

**Figure S5. Lip2 esterifies AFAs with cholesterol.**

**A**, Thin layer chromatography of lipids extracted after incubation of PA, SA, OA (oleic acid), LA, or ALA (α-linoleic acid) with Chol in the presence of recombinant *Staphylococcus aureus* lipase 2 (Lip2) or catalytically dead Lip2 S412A. Cholesteryl esters (CE) were detected for all AFAs tested. **B**, TLC lipid analysis of USA300 *S. aureus*-conditioned media from the indicated strain (WT pEmpty, Δlip pEmpty, Δlip p*lip1*, Δlip p*lip2*, or Δlip p*lip2^S^*^412^*^A^*) incubated with Chol and ALA. **C**, UHPLC-MS/MS lipid analysis to measure cholesterol upon incubation of *S. aureus*-conditioned media from strains described in **B** with Chol and LA. **D**, TLC of lipids extracted after incubation of *S. aureus*-conditioned media from strains listed in **B** with Chol, ethanol, and ALA. Ethyl esters (EE) and/or CE were detected. **E-F**, TLC lipid analysis of *S. aureus*-conditioned media from wild-type USA400 MW2 (MW2 WT), or its lipase-deficient mutants (MW2 Δ*lip1* and MW2 Δ*lip2*) incubated with Chol and ALA in the absence (**E**) or presence of ethanol (**F**). Four lipid standards (ALA, cholesterol, cholesteryl ALA, and ethyl ALA are shown. Bar graphs (**C**) are means + SEM for five biological replicates. Statistical significance by one-way ANOVA with Dunnett’s test relative to WT pEmpty. \*\*\**P* < 0.001, \*\*\*\**P* < 0.0001.

**Figure S6. Cholesterol does not prevent membrane-damaging effects of AFAs.**

**A**, Wild-type USA300 JE2 and its isogenic Δlip mutant with pEmpty, and Δlip complemented with p*lip1*, p*lip2^S^*^412^*^A^*, or p*lip2* were left untreated or treated with dehydroergosterol. After washing with PBS, DHE-binding was quantified by fluorometry in relative fluorescence units (RFU). **B-C**, WT pEmpty, Δlip pEmpty, and Δlip p*lip2* were stained with azide fluor 488 upon incubation in plain NB, or NB supplemented with palmitoleic acid (PA) alkyne or PA alkyne + cholesterol. RFU or mean fluorescence intensities (MFI) were determined using fluorometry (**B**) or flow cytometry (**C**), respectively. **D**, The indicated strain (WT pEmpty, Δlip pEmpty, Δlip p*lip1*, or Δlip p*lip2*) was incubated in NB, or NB supplemented with PA or PA + Chol prior to staining with DiOC_2_(3) (3,3′-diethyloxacarbocyanine iodide). Membrane potential, as computed by the ratio between red and green fluorescence intensities (“red shift”), was determined by fluorometry. Shown are mean + SEM for three (**A-C**) or four (**D**) biological replicates. Two-way ANOVA with Tukey’s multiple comparisons test was used to calculate statistical significance (**B**). \**P* < 0.05, \*\**P* < 0.01, \*\*\**P* < 0.001, \*\*\*\**P* < 0.0001.

**Fig. S7. Lip2 is conserved in *S. aureus*.**

Lip2 is generally synthesized as a 690 or 691 amino acid polypeptide. A consensus Lip2 sequence was generated upon alignment of over 3000 Lip2 sequences from our database to USA300 Lip2 as reference. The percentage of the modal residue at each amino acid position is shown.

**Fig. S8. The sequence type dictates Lip2 diversity.**

The multiple sequence alignment of over 3000 Lip2 sequences is represented as three-dimensional space generated using dimensionality reduction. Lip2 sequence of each *S. aureus* strain is represented as a dot whose colour depends either on the isolation site (**A**) or the sequence type (ST) (**B**) of the bacterium.

**Fig. S9. Lip2 displays mutation hotspots and is disproportionally disrupted in ST398 strains.**

**A**, Lip2 is usually a 690 or 691 amino acid protein. For the > 3000 Lip2 sequences from our database, the mutation rate at each amino acid position, relative to USA300, was determined. The insertion of serine (S) between positions 43 and 44 in ∼ 70% of our strains is denoted as “-44S” and highlighted in red as well as all mutations that occurred in at least a quarter of our database. **B-C**, Sequence types (ST) of all *S. aureus* isolates in our database (**B**) or isolates with prophage-disrupted Lip2.

## Notes

### Competing Interest Statement

The authors have declared no competing interest.

## References

1 Hines, K. M. et al. Lipidomic and Ultrastructural Characterization of the Cell Envelope of Staphylococcus aureus Grown in the Presence of Human Serum. mSphere 5 (2020). https://doi.org:10.1128/mSphere.00339-20

2 Frank, M. W. et al. Host Fatty Acid Utilization by Staphylococcus aureus at the Infection Site. mBio 11 (2020). https://doi.org:10.1128/mBio.00920-20

3 Feingold, K. R. & Elias, P. M. Role of lipids in the formation and maintenance of the cutaneous permeability barrier. Biochim Biophys Acta 1841, 280–294 (2014). https://doi.org:10.1016/j.bbalip.2013.11.007

4 Zheng, Y. et al. Commensal Staphylococcus epidermidis contributes to skin barrier homeostasis by generating protective ceramides. Cell Host Microbe 30, 301–313.e309 (2022). https://doi.org:10.1016/j.chom.2022.01.004

5 Rivera-Chavez, F. & Mekalanos, J. J. Cholera toxin promotes pathogen acquisition of host-derived nutrients. Nature 572, 244–248 (2019). https://doi.org:10.1038/s41586-019-1453-3

6 Eierhoff, T. et al. A lipid zipper triggers bacterial invasion. Proc Natl Acad Sci U S A 111, 12895–12900 (2014). https://doi.org:10.1073/pnas.1402637111

7 Bae, M. et al. Akkermansia muciniphila phospholipid induces homeostatic immune responses. Nature 608, 168–173 (2022). https://doi.org:10.1038/s41586-022-04985-7

8 Clarke, S. R. et al. The Staphylococcus aureus surface protein IsdA mediates resistance to innate defenses of human skin. Cell Host Microbe 1, 199–212 (2007). https://doi.org:10.1016/j.chom.2007.04.005

9 Do, T. Q. et al. Lipids including cholesteryl linoleate and cholesteryl arachidonate contribute to the inherent antibacterial activity of human nasal fluid. J Immunol 181, 4177–4187 (2008). https://doi.org:10.4049/jimmunol.181.6.4177

10 Verhaegh, R., Becker, K. A., Edwards, M. J. & Gulbins, E. Sphingosine kills bacteria by binding to cardiolipin. J Biol Chem 295, 7686–7696 (2020). https://doi.org:10.1074/jbc.RA119.012325

11 Flores-Díaz, M., Monturiol-Gross, L., Naylor, C., Alape-Girón, A. & Flieger, A. Bacterial Sphingomyelinases and Phospholipases as Virulence Factors. Microbiol Mol Biol Rev 80, 597–628 (2016). https://doi.org:10.1128/mmbr.00082-15

12 Bomar, L., Brugger, S. D., Yost, B. H., Davies, S. S. & Lemon, K. P. Corynebacterium accolens Releases Antipneumococcal Free Fatty Acids from Human Nostril and Skin Surface Triacylglycerols. mBio 7, e01725–01715 (2016). https://doi.org:10.1128/mBio.01725-15

13 Chen, X. & Alonzo, F, 3rd.., Bacterial lipolysis of immune-activating ligands promotes evasion of innate defenses. Proc Natl Acad Sci U S A 116, 3764–3773 (2019). https://doi.org:10.1073/pnas.1817248116

14 Kengmo Tchoupa, A., Eijkelkamp, B. A. & Peschel, A. Bacterial adaptation strategies to host-derived fatty acids. Trends Microbiol 30, 241–253 (2022). https://doi.org:10.1016/j.tim.2021.06.002

15 Krismer, B., Weidenmaier, C., Zipperer, A. & Peschel, A. The commensal lifestyle of Staphylococcus aureus and its interactions with the nasal microbiota. Nature reviews. Microbiology 15, 675–687 (2017). https://doi.org:10.1038/nrmicro.2017.104

16 Williams, M. R. & Gallo, R. L. The role of the skin microbiome in atopic dermatitis. Curr Allergy Asthma Rep 15, 65 (2015). https://doi.org:10.1007/s11882-015-0567-4

17 Takigawa, H., Nakagawa, H., Kuzukawa, M., Mori, H. & Imokawa, G. Deficient production of hexadecenoic acid in the skin is associated in part with the vulnerability of atopic dermatitis patients to colonization by Staphylococcus aureus. Dermatology 211, 240–248 (2005). https://doi.org:10.1159/000087018

18 Mortensen, J. E., Shryock, T. R. & Kapral, F. A. Modification of bactericidal fatty acids by an enzyme of Staphylococcus aureus. J Med Microbiol 36, 293–298 (1992). https://doi.org:10.1099/00222615-36-4-293

19 Neumann, Y. et al. The effect of skin fatty acids on Staphylococcus aureus. Arch Microbiol 197, 245-267 (2015). https://doi.org:10.1007/s00203-014-1048-1

20 Kengmo Tchoupa, A. & Peschel, A. Staphylococcus aureus Releases Proinflammatory Membrane Vesicles To Resist Antimicrobial Fatty Acids. mSphere 5 (2020). https://doi.org:10.1128/mSphere.00804-20

21 Nguyen, M. T., Hanzelmann, D., Härtner, T., Peschel, A. & Götz, F. Skin-Specific Unsaturated Fatty Acids Boost the Staphylococcus aureus Innate Immune Response. Infect Immun 84, 205–215 (2016). https://doi.org:10.1128/iai.00822-15

22 Subramanian, C., Frank, M. W., Batte, J. L., Whaley, S. G. & Rock, C. O. Oleate hydratase from Staphylococcus aureus protects against palmitoleic acid, the major antimicrobial fatty acid produced by mammalian skin. J Biol Chem 294, 9285–9294 (2019). https://doi.org:10.1074/jbc.RA119.008439

23 Cadieux, B., Vijayakumaran, V., Bernards, M. A., McGavin, M. J. & Heinrichs, D. E. Role of lipase from community-associated methicillin-resistant Staphylococcus aureus strain USA300 in hydrolyzing triglycerides into growth-inhibitory free fatty acids. J Bacteriol 196, 4044–4056 (2014). https://doi.org:10.1128/JB.02044-14

24 Bae, T., Baba, T., Hiramatsu, K. & Schneewind, O. Prophages of Staphylococcus aureus Newman and their contribution to virulence. Mol Microbiol 62, 1035–1047 (2006). https://doi.org:10.1111/j.1365-2958.2006.05441.x

25 Zorn, K., Oroz-Guinea, I., Brundiek, H. & Bornscheuer, U. T. Engineering and application of enzymes for lipid modification, an update. Prog Lipid Res 63, 153–164 (2016). https://doi.org:10.1016/j.plipres.2016.06.001

26 Chimalapati, S. et al. Vibrio deploys type 2 secreted lipase to esterify cholesterol with host fatty acids and mediate cell egress. Elife 9 (2020). https://doi.org:10.7554/eLife.58057

27 Long, J. P., Hart, J., Albers, W. & Kapral, F. A. The production of fatty acid modifying enzyme (FAME) and lipase by various staphylococcal species. J Med Microbiol 37, 232–234 (1992). https://doi.org:10.1099/00222615-37-4-232

28 Pourmousa, M. et al. Dehydroergosterol as an analogue for cholesterol: why it mimics cholesterol so well-or does it? J Phys Chem B 118, 7345–7357 (2014). https://doi.org:10.1021/jp406883k

29 Kengmo Tchoupa, A., et al. The type VII secretion system protects Staphylococcus aureus against antimicrobial host fatty acids. Sci Rep 10, 14838 (2020). https://doi.org:10.1038/s41598-020-71653-z

30 Kenny, J. G. et al. The Staphylococcus aureus response to unsaturated long chain free fatty acids: survival mechanisms and virulence implications. PLoS One 4, e4344 (2009). https://doi.org:10.1371/journal.pone.0004344

31 Cartron, M. L. et al. Bactericidal activity of the human skin fatty acid cis-6-hexadecanoic acid on Staphylococcus aureus. Antimicrob Agents Chemother 58, 3599–3609 (2014). https://doi.org:10.1128/AAC.01043-13

32 Davis, J. J. et al. The PATRIC Bioinformatics Resource Center: expanding data and analysis capabilities. Nucleic Acids Res 48, D606–D612 (2020). https://doi.org:10.1093/nar/gkz943

33 Ozer, E. A., Nnah, E., Didelot, X., Whitaker, R. J. & Hauser, A. R. The Population Structure of Pseudomonas aeruginosa Is Characterized by Genetic Isolation of exoU+ and exoS+ Lineages. Genome Biol Evol 11, 1780–1796 (2019). https://doi.org:10.1093/gbe/evz119

34 Nakamura, K., Williams, M. R., Kwiecinski, J. M., Horswill, A. R. & Gallo, R. L. Staphylococcus aureus Enters Hair Follicles Using Triacylglycerol Lipases Preserved through the Genus Staphylococcus. J Invest Dermatol 141, 2094–2097 (2021). https://doi.org:10.1016/j.jid.2021.02.009

35 Zipperer, A. et al. Human commensals producing a novel antibiotic impair pathogen colonization. Nature 535, 511–516 (2016). https://doi.org:10.1038/nature18634

36 Nguyen, M. T. et al. Staphylococcal (phospho)lipases promote biofilm formation and host cell invasion. Int J Med Microbiol 308, 653–663 (2018). https://doi.org:10.1016/j.ijmm.2017.11.013

37 Nguyen, M. T., Matsuo, M., Niemann, S., Herrmann, M. & Götz, F. Lipoproteins in Gram-Positive Bacteria: Abundance, Function, Fitness. Front Microbiol 11, 582582 (2020). https://doi.org:10.3389/fmicb.2020.582582

38 Kim, B. E. et al. Side-by-Side Comparison of Skin Biopsies and Skin Tape Stripping Highlights Abnormal Stratum Corneum in Atopic Dermatitis. J Invest Dermatol 139, 2387–2389 e2381 (2019). https://doi.org:10.1016/j.jid.2019.03.1160

39 Li, S. et al. Altered composition of epidermal lipids correlates with Staphylococcus aureus colonization status in atopic dermatitis. Br J Dermatol 177, e125–e127 (2017). https://doi.org:10.1111/bjd.15409

40 Gerlach, D. et al. Methicillin-resistant Staphylococcus aureus alters cell wall glycosylation to evade immunity. Nature 563, 705–709 (2018). https://doi.org:10.1038/s41586-018-0730-x

41 Kamal, F. et al. Differential Cellular Response to Translocated Toxic Effectors and Physical Penetration by the Type VI Secretion System. Cell Rep 31, 107766 (2020). https://doi.org:10.1016/j.celrep.2020.107766

42 Deruelle, V. et al. The bacterial toxin ExoU requires a host trafficking chaperone for transportation and to induce necrosis. Nat Commun 12, 4024 (2021). https://doi.org:10.1038/s41467-021-24337-9

43 Urbanek, A. et al. Composition and antimicrobial activity of fatty acids detected in the hygroscopic secretion collected from the secretory setae of larvae of the biting midge Forcipomyia nigra (Diptera: Ceratopogonidae). J Insect Physiol 58, 1265–1276 (2012). https://doi.org:10.1016/j.jinsphys.2012.06.014

44 Rosenstein, R. & Gotz, F. Staphylococcal lipases: biochemical and molecular characterization. Biochimie 82, 1005–1014 (2000). https://doi.org:10.1016/s0300-9084(00)01180-9

45 Christensson, B., Fehrenbach, F. J. & Hedstrom, S. A. A new serological assay for Staphylococcus aureus infections: detection of IgG antibodies to S. aureus lipase with an enzyme-linked immunosorbent assay. J Infect Dis 152, 286–292 (1985). https://doi.org:10.1093/infdis/152.2.286

46 Lowe, A. M., Beattie, D. T. & Deresiewicz, R. L. Identification of novel staphylococcal virulence genes by in vivo expression technology. Mol Microbiol 27, 967–976 (1998). https://doi.org:10.1046/j.1365-2958.1998.00741.x

47 Hu, C., Xiong, N., Zhang, Y., Rayner, S. & Chen, S. Functional characterization of lipase in the pathogenesis of Staphylococcus aureus. Biochem Biophys Res Commun 419, 617–620 (2012). https://doi.org:10.1016/j.bbrc.2012.02.057

48 Ishii, K. et al. Induction of virulence gene expression in Staphylococcus aureus by pulmonary surfactant. Infect Immun 82, 1500–1510 (2014). https://doi.org:10.1128/IAI.01635-13

49 Cheung, G. Y., Wang, R., Khan, B. A., Sturdevant, D. E. & Otto, M. Role of the accessory gene regulator agr in community-associated methicillin-resistant Staphylococcus aureus pathogenesis. Infect Immun 79, 1927–1935 (2011). https://doi.org:10.1128/IAI.00046-11

50 Blevins, J. S., Beenken, K. E., Elasri, M. O., Hurlburt, B. K. & Smeltzer, M. S. Strain-dependent differences in the regulatory roles of sarA and agr in Staphylococcus aureus. Infect Immun 70, 470–480 (2002). https://doi.org:10.1128/IAI.70.2.470-480.2002

51 Jones, R. C., Deck, J., Edmondson, R. D. & Hart, M. E. Relative quantitative comparisons of the extracellular protein profiles of Staphylococcus aureus UAMS-1 and its sarA, agr, and sarA agr regulatory mutants using one-dimensional polyacrylamide gel electrophoresis and nanocapillary liquid chromatography coupled with tandem mass spectrometry. J Bacteriol 190, 5265–5278 (2008). https://doi.org:10.1128/JB.00383-08

52 Chamberlain, N. R. & Imanoel, B. Genetic regulation of fatty acid modifying enzyme from Staphylococcus aureus. J Med Microbiol 44, 125–129 (1996). https://doi.org:10.1099/00222615-44-2-125

53 Gajenthra Kumar, N., et al. Untargeted lipidomic analysis to broadly characterize the effects of pathogenic and non-pathogenic staphylococci on mammalian lipids. PLoS One 13, e0206606 (2018). https://doi.org:10.1371/journal.pone.0206606

54 Monk, I. R., Shah, I. M., Xu, M., Tan, M. W. & Foster, T. J. Transforming the untransformable: application of direct transformation to manipulate genetically Staphylococcus aureus and Staphylococcus epidermidis. mBio 3 (2012). https://doi.org:10.1128/mBio.00277-11

55 Bateman, B. T., Donegan, N. P., Jarry, T. M., Palma, M. & Cheung, A. L. Evaluation of a tetracycline-inducible promoter in Staphylococcus aureus in vitro and in vivo and its application in demonstrating the role of sigB in microcolony formation. Infect Immun 69, 7851–7857 (2001). https://doi.org:10.1128/IAI.69.12.7851-7857.2001

56 Schlatterer, K. et al. The Mechanism behind Bacterial Lipoprotein Release: Phenol-Soluble Modulins Mediate Toll-Like Receptor 2 Activation via Extracellular Vesicle Release from Staphylococcus aureus. mBio 9 (2018). https://doi.org:10.1128/mBio.01851-18

57 Cheung, G. Y. et al. Functional characteristics of the Staphylococcus aureus delta-toxin allelic variant G10S. Sci Rep 5, 18023 (2015). https://doi.org:10.1038/srep18023

58 Bligh, E. G. & Dyer, W. J. A rapid method of total lipid extraction and purification. Can J Biochem Physiol 37, 911–917 (1959). https://doi.org:10.1139/o59-099

59 Matyash, V., Liebisch, G., Kurzchalia, T. V., Shevchenko, A. & Schwudke, D. Lipid extraction by methyl-tert-butyl ether for high-throughput lipidomics. J Lipid Res 49, 1137–1146 (2008). https://doi.org:10.1194/jlr.D700041-JLR200

60 Huang, L. et al. Molecular Basis of Rhodomyrtone Resistance in Staphylococcus aureus. mBio 13, e0383321 (2022). https://doi.org:10.1128/mbio.03833-21

61 Arndt, D. et al. PHASTER: a better, faster version of the PHAST phage search tool. Nucleic Acids Res 44, W16–21 (2016). https://doi.org:10.1093/nar/gkw387

62 Katoh, K. & Standley, D. M. MAFFT multiple sequence alignment software version 7: improvements in performance and usability. Mol Biol Evol 30, 772–780 (2013). https://doi.org:10.1093/molbev/mst010

