## Supplementary figures and images for "Lipase-mediated detoxification of host-derived antimicrobial fatty acids by *Staphylococcus aureus*"

### Supplemental Figure 1

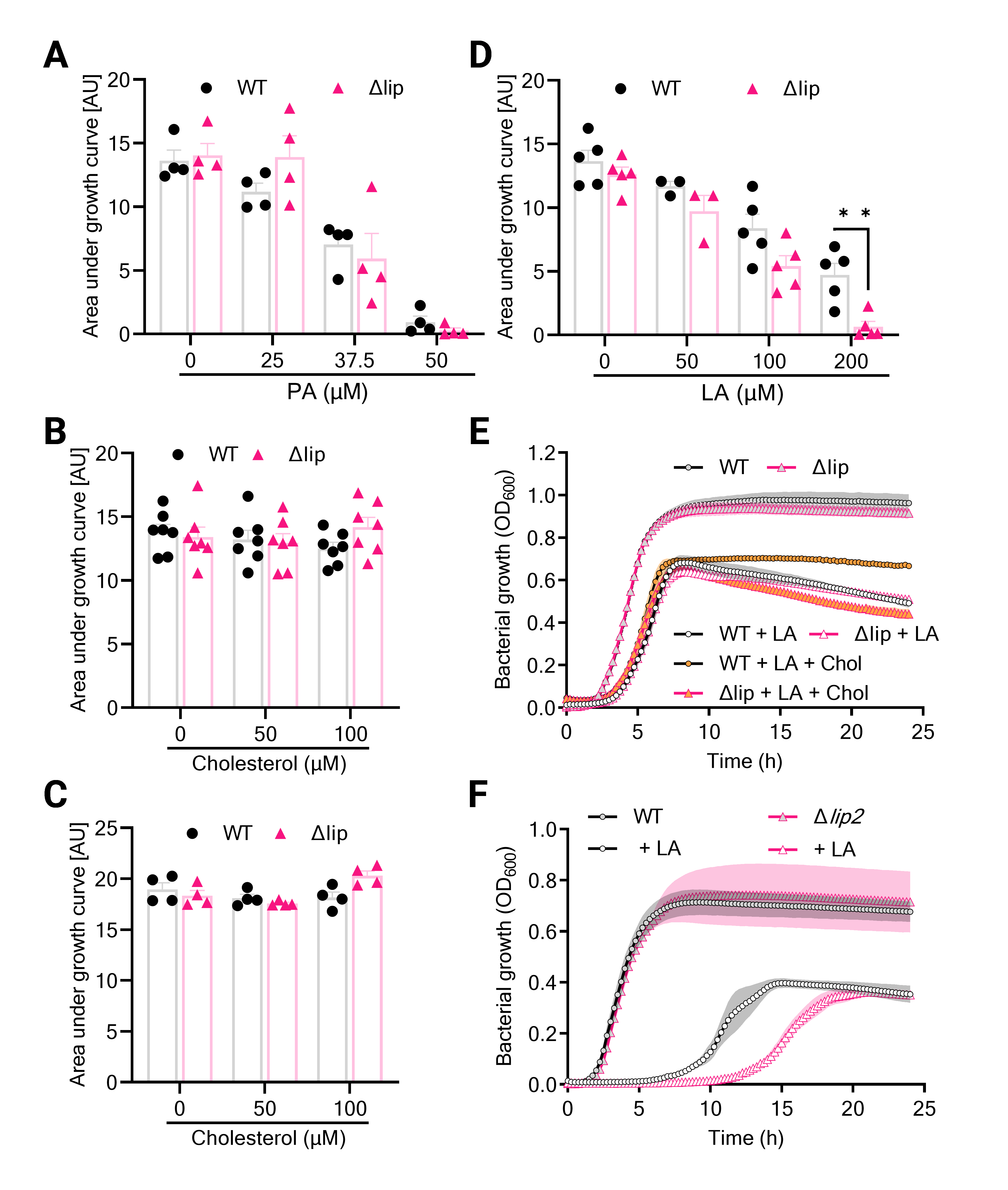

### Supplemental Figure 2

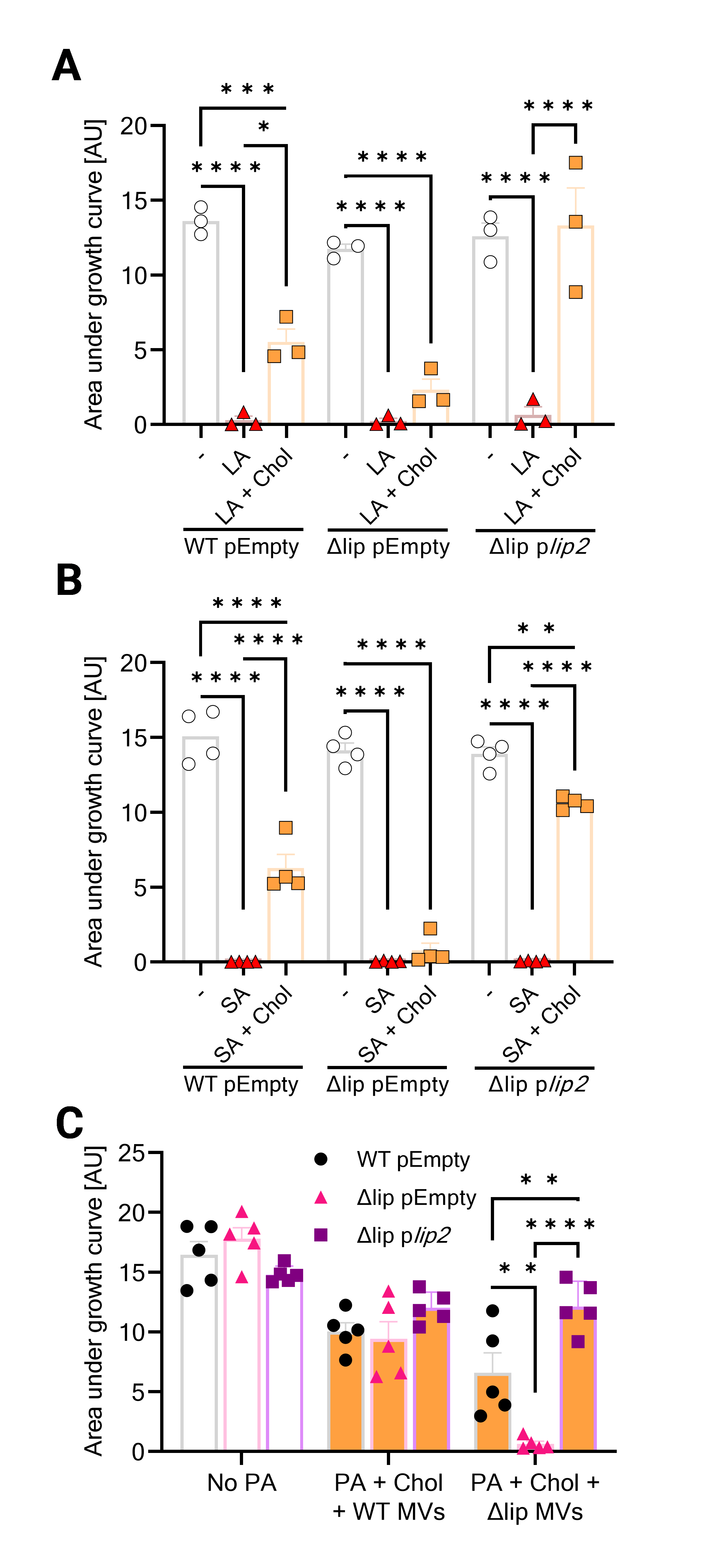

### Supplemental Figure 3

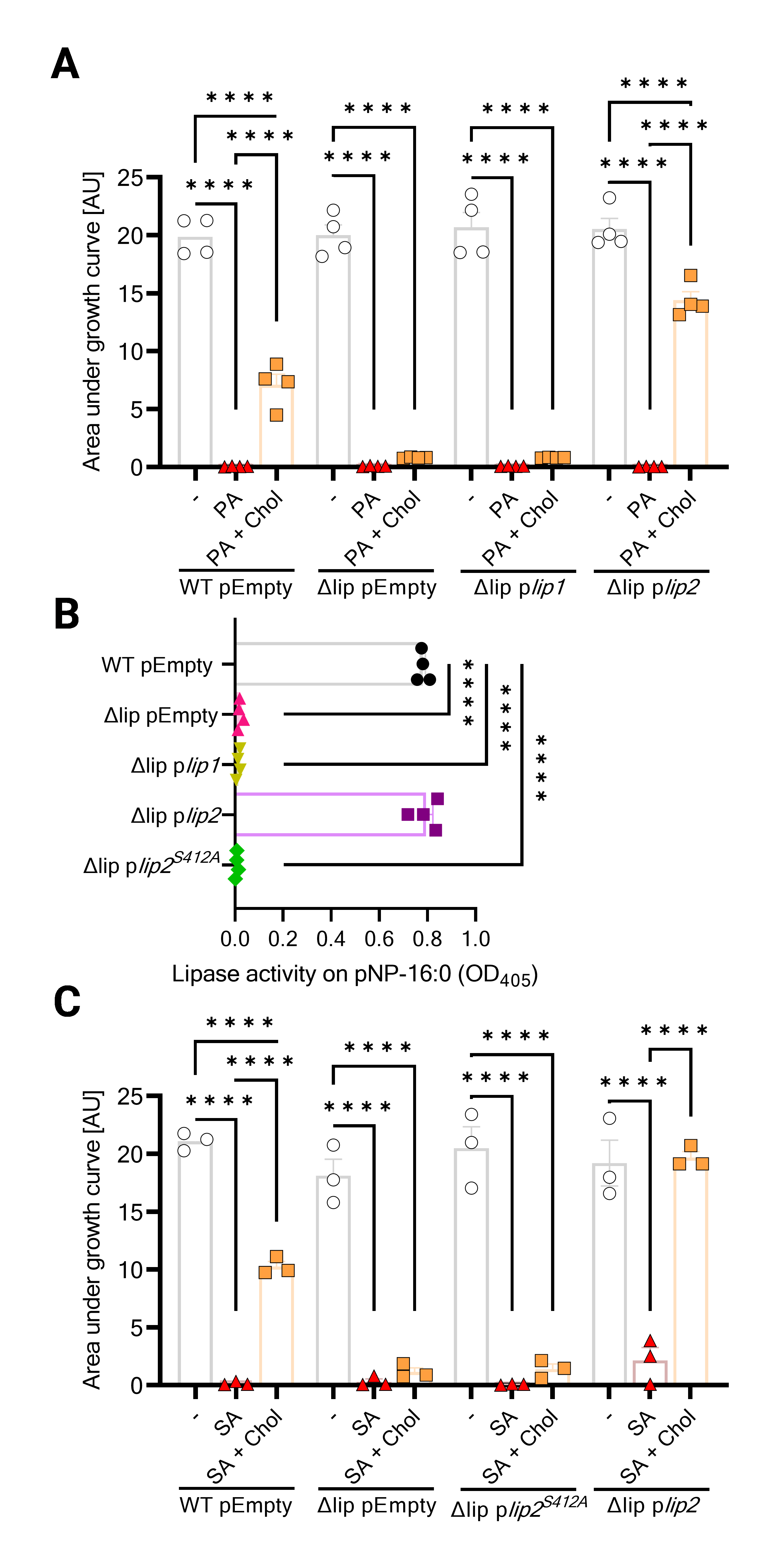

### Supplemental Figure 4

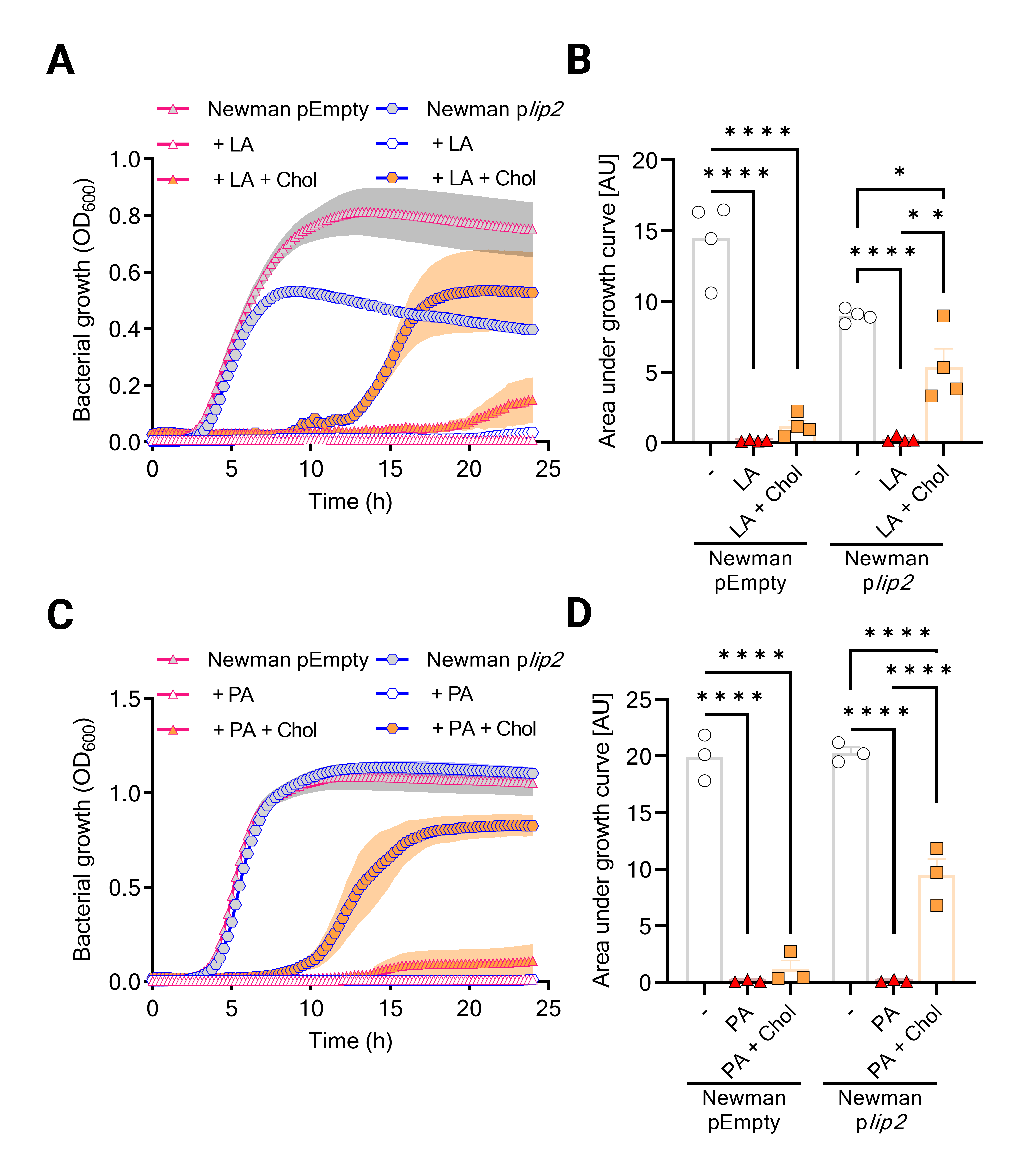

### Supplemental Figure 5

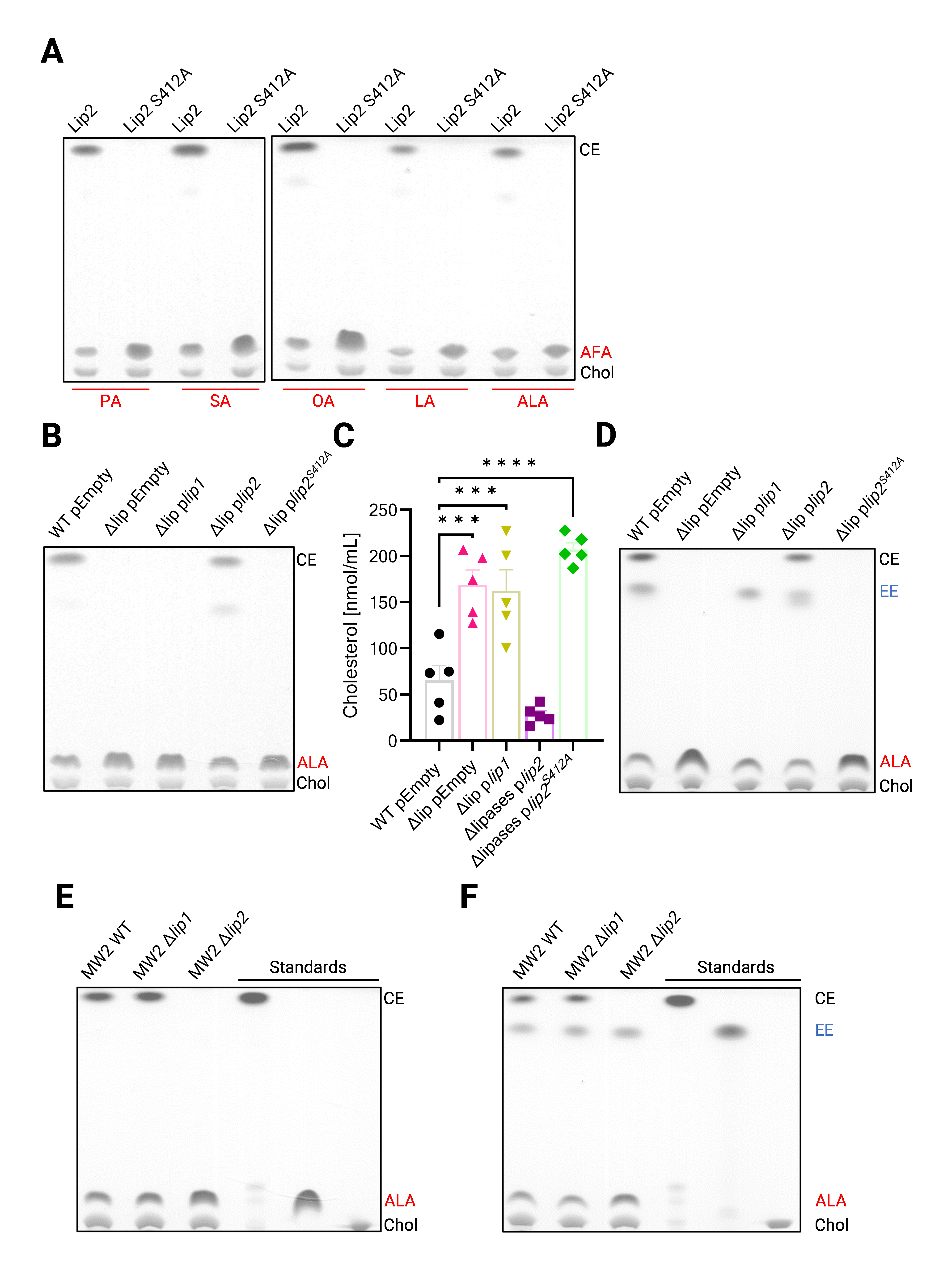

### Supplemental Figure 6

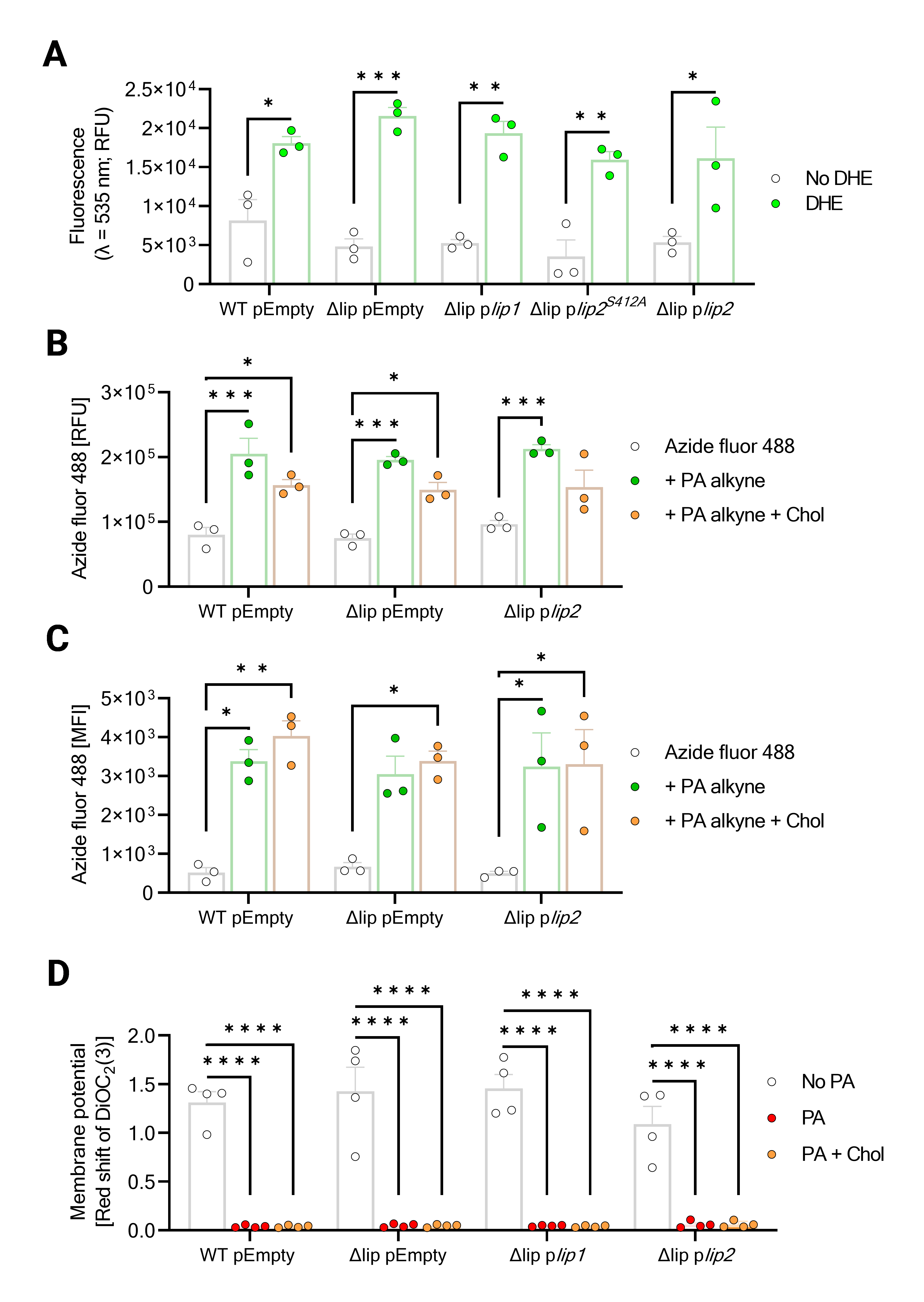

### Supplemental Figure 7

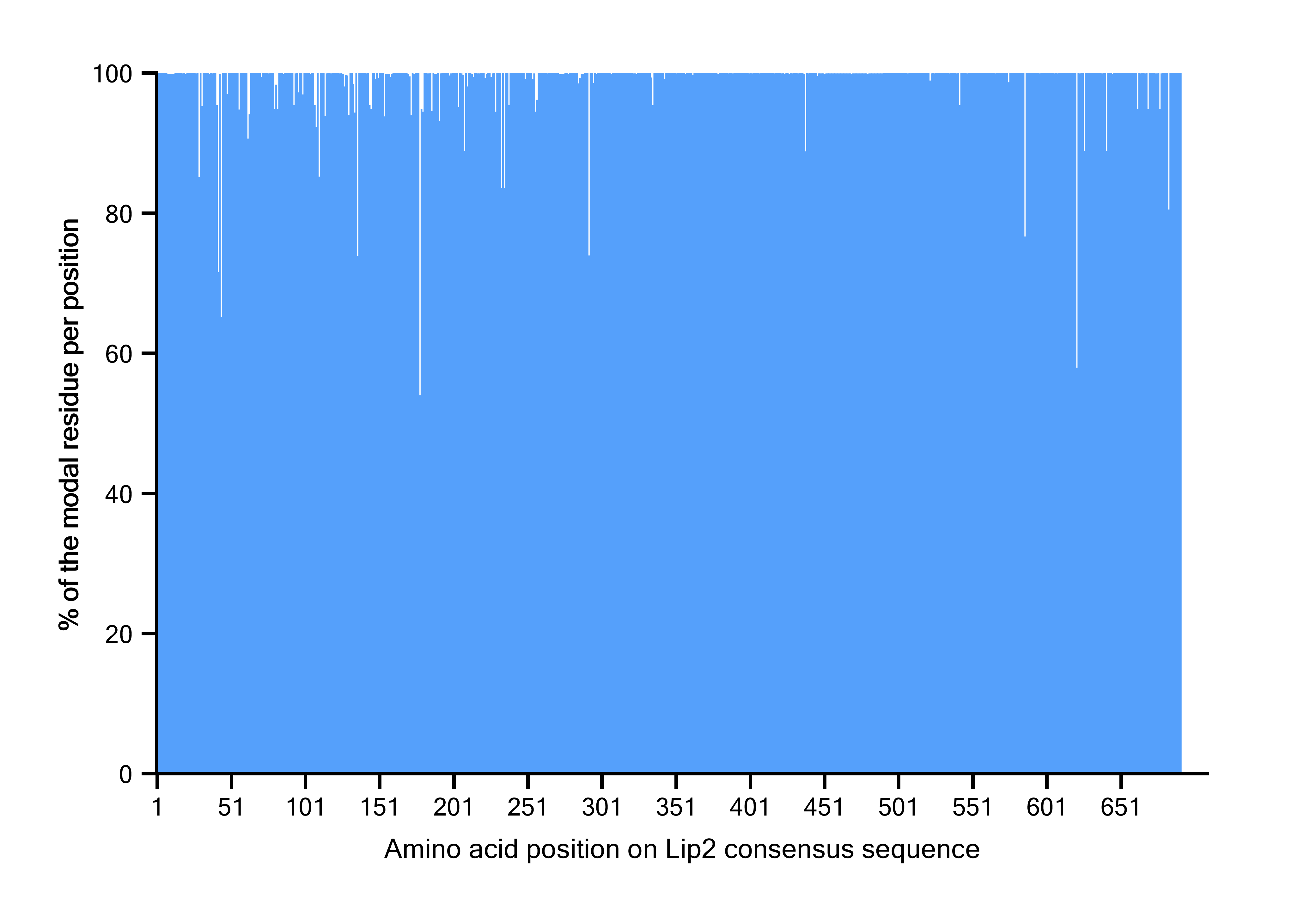

### Supplemental Figure 8

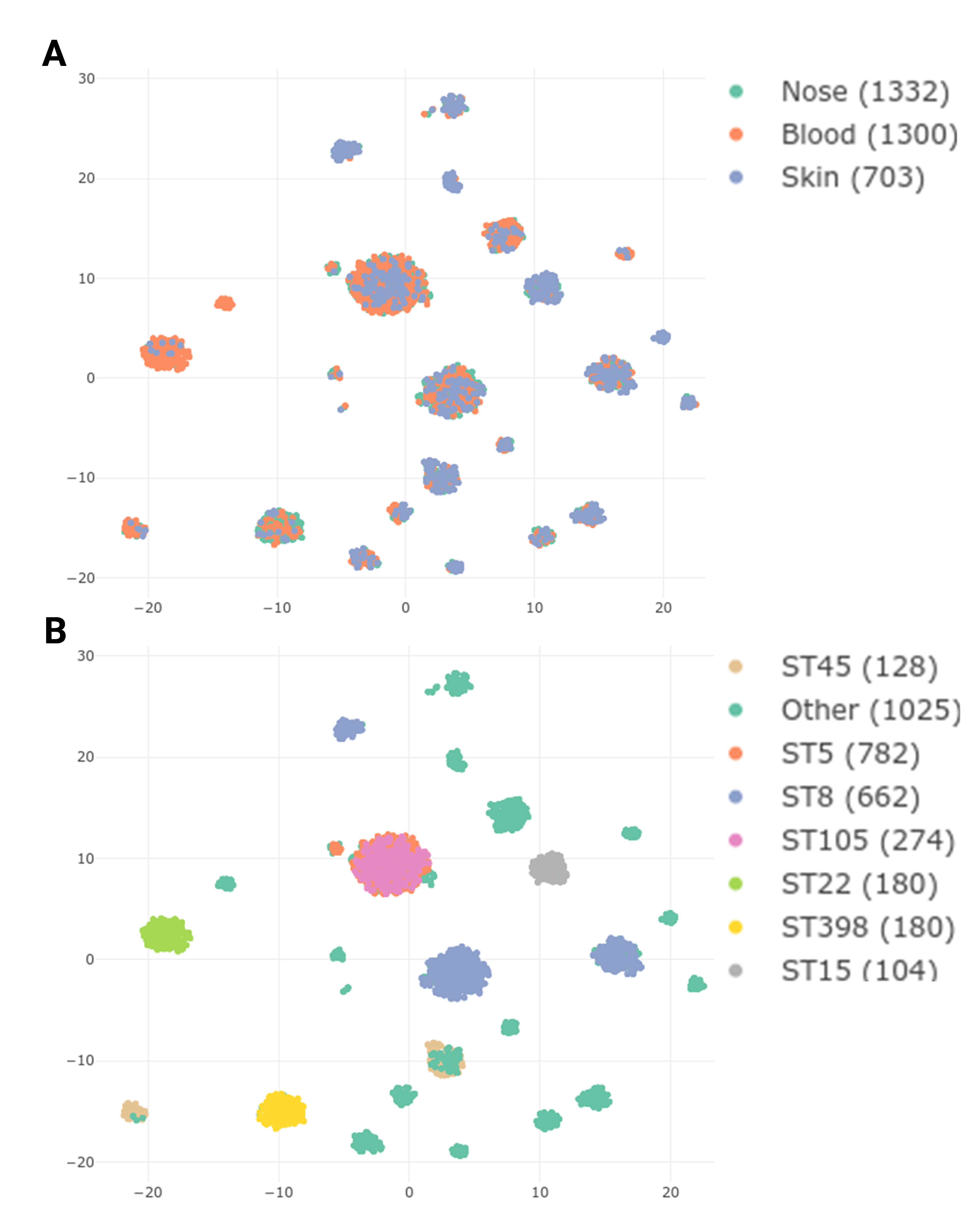

### Supplemental Figure 9

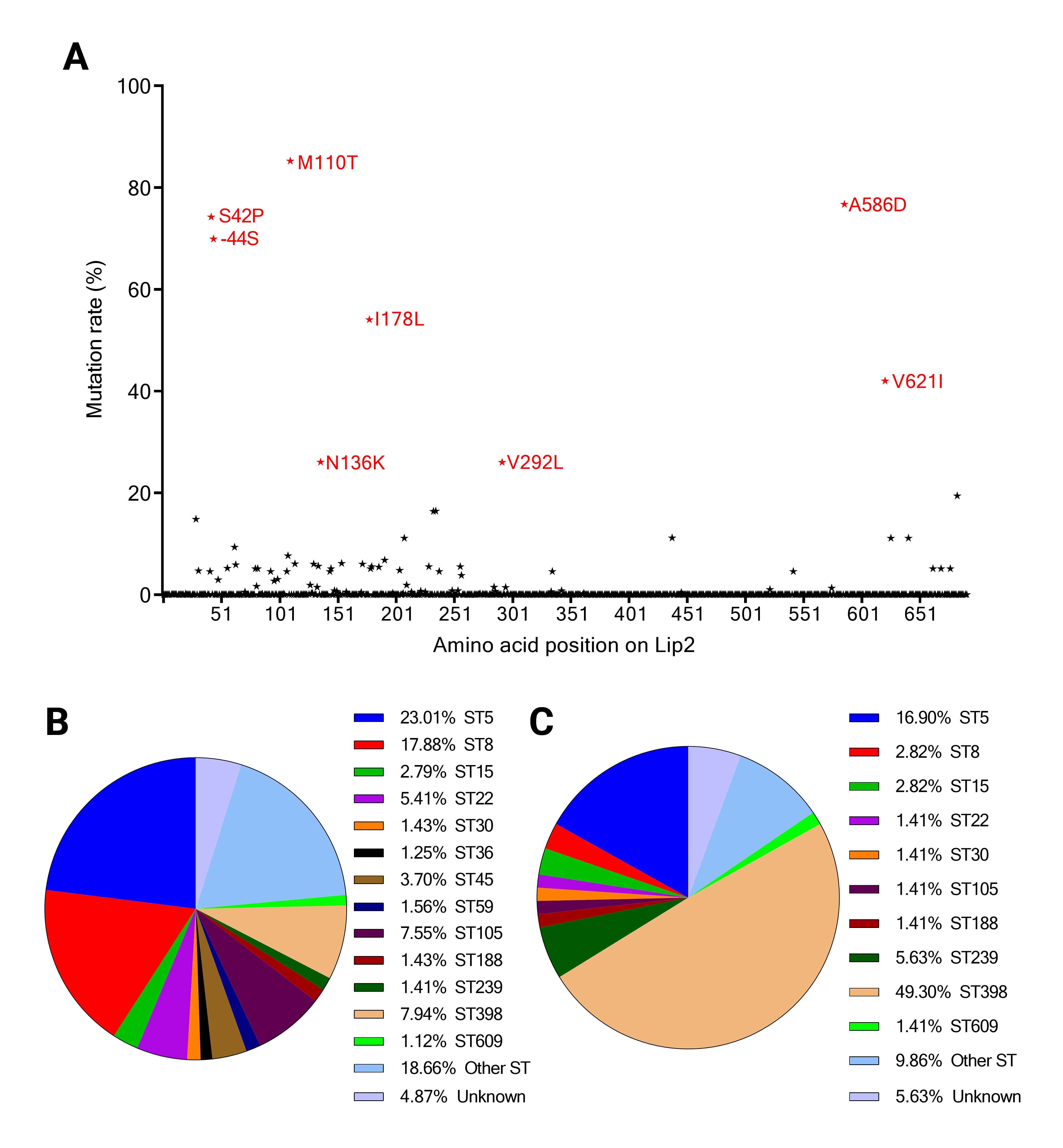
