## Supplemental Table 1 for "Lipase-mediated detoxification of host-derived antimicrobial fatty acids by *Staphylococcus aureus*"

**Table S1: Mutations co-occurring in Lip2**

| Mutations | Strain number |
|---|---|
| S42P,-44S,M110T,I178L,A586D,V621I | 1174 |
| M110T | 199 |
| V29I,S42P,-44S,I63V,I108S,M110T,Q130P,N136K,G154E,I172V,P191L,T208S,T233N,R235H,V292I, K438Q,A586D,D626N,I641L,S683G, | 190 |
| V29I,S42P,-44S,A62T,M110T,T134I,N136K,I178L,N180H,K229T,P256S,V292I,A586D,V621I,S683G | 181 |
| S31T,T41I,S42P,-44T,T93M,T107I,M110T,N136K,V144I,A204E,T233N,R235H,E238N,F335Y D542N,A586D | 145 |
| S42P,-44S,A62V,M110T,I178L,E257D,V292I,A586D | 125 |
| S42P,-44S,P56S,E80K,G82D,M110T,N114D,N136K,Q145L,P179T,P186T,T208S,T233N,R235H,V292I, K438Q, A586D,D626N,I641L,A662S,T669I,R677S,S683G | 105 |
| S42P,A48V,S96L,H99Y,M110T,I178L,A586D | 66 |
| S42P,-44S,M110T,Q127H,I178L,D210E,A586D | 61 |
| V29I,S42L,-44S,I108S,M110T,N136K,V292I,A586D,S683G | 57 |
| N81D,M110T | 54 |
| M110T,A586D | 48 |
| S42P,-44S,P56S,E80K,G82D,M110T,N114D,N136K,Q145L,P179T,P186T,T208S,T233N,R235H,A285-, V292I,K438Q,A586D,D626N,I641L,A662S,T669I,R677S,S683G | 46 |
| M110T,R522H,A586D | 32 |
| S42P,-44S,M110T,A133V,I178L,V292I,L295I,A586D | 32 |
| S42P,M110T,N136K,P191T,K343N,A586D | 27 |
| V29I,S42L,-44S,M110T,N114D,N136K,H148R,H249R,N254K,V292I,A586D,S683G | 25 |
| S42P,M110T,N136K,A222V,T233N,R235H,A286T,V292I,A586D | 23 |
| S42P,A48V,S96L,H99Y,M110T,I178L,K334N,A586D | 21 |
| S42P,-44S,M110T,I178L,E575A,A586D | 18 |
| V29I,S42P,-44S,G71R,M110T,N136K,A150T,I178L,A586D | 17 |
| S42P,-44S,M110T,T158R,S171F,I178L,I226T,V292I,A586D | 15 |
| S42P,-44S,M110T,I178L,N214S,A586D,V621I | 14 |
| S42P,-44S,P56S,E80K,G82D,M110T,N114D,N136K,Q145L,P179T,P186T,T208S,T233N,R235H,V292I, K438Q,P446S,A586D,D626N,I641L,A662S,T669I,R677S,S683G | 13 |
| M110T,E575G | 12 |
| S42P,-44S,H99Q,M110T,H129N,G154R,I178L,P186T,V292I,A586D | 10 |
| S42P,-44S,M110T,A133V,I178L,A198E,V292I,L295I,R362C,A586D | 8 |
| M110T,A207E | 6 |
| S42P,-44S,M110T,S128P,I178L,A586D,V621I | 6 |
| S42P,-44S,M110T,I178L,A297E,E575A,A586D | 5 |
| S42P,-44S,M110T,A133V,I178L,H223Y,V292I,L295I,A586D | 4 |
| V29I,S42P,-44S,T86I,M110T,N114D,Q130P,N136K,I172V,T208S,T233N,R235H,V292I,K438Q,A586D, D626N,I641L,S683G | 4 |
| M110T,D272-,A273-,L274-,Q275- | 4 |
| S42P,-44S,N104I,M110T,I178L,A586D,V621I | 4 |
| S42P,-44S,M110T,A133V,I178L,V292I,L295I,A379V,A586D | 4 |
| V20L,V29I,S42P,-44S,A48T,M110T,A150T,S188P,P191T,K278N,A586D | 4 |
| S42P,A48V,S96L,H99Y,M110T,I178L,A204V,A586D | 4 |
| S42P,-44S,M110T,I178L,N214S,A324S,A586D,V621I | 3 |
| M110T,A174T | 3 |
| R8-,K9-,Y10-,S11-,I12-,S42P,-44S,M110T,I178L,E575A,A586D | 3 |
