## Supplemental Table 2 for "Lipase-mediated detoxification of host-derived antimicrobial fatty acids by *Staphylococcus aureus*"

**Table S2. Strains and plasmids**

| Strain or plasmid | Description | Source or reference |
|---|---|---|
| ***Staphylococcus aureus*** | | |
| Newman | Methicillin-sensitive *S. aureus* (MSSA) | [1] |
| SH1000 | NCTC8325 derivative with a functional *rsbU* gene, Δ*tcaR*, cured of φ11, φ12, and φ13; MSSA | Simon Foster [2] |
| USA300 | Community-acquired MRSA (CA-MRSA), plasmid-cured derivative of LAC strain | David E. Heinrichs [3] |
| USA300 Δ*lip2* | USA300 with the *lip2* (*gehB*) gene deleted | David E. Heinrichs [3] |
| USA300 JE2 | CA-MRSA, plasmid-cured derivative of LAC strain | Paul Fey [4] |
| USA300 JE2 Δlip | USA300 JE2 defective for Lip1 and Lip2 | Friedrich Götz [5] |
| Newman | Methicillin-sensitive *S. aureus* (MSSA) | [1] |
| USA400 MW2 | CA-MRSA | Michael Otto [6] |
| USA400 MW2 Δ*lip1* | MW2 with the *lip1* (*gehA*) gene deleted | This study |
| USA400 MW2 Δ*lip2* | MW2 with the *lip2* (*gehB*) gene deleted | This study |
| ***Escherichia coli*** | | |
| BL21 (DE3) | $F^-$ *ompT hsdS*$_B$($r_B^-$ $m_B^-$) *gal dcm* (DE3), IPTG-inducible T7 RNA polymerase | |
| DC10B | Δ*dcm* in the DH10B background (K-12 derivative) | Simon Heilbronner [7] |
| SA08B | DC10BΩ$P_{help}$-*hsdMS* (CC8-2) of NRS384 integrated between the *atpI* and *gidB* genes | Simon Heilbronner [8] |
| IM01B | SA08BΩ$P_{N25}$-*hsdS* (CC1-1) of MW2 integrated between the *essQ* and *cspB* genes | Simon Heilbronner [8] |
| **Plasmid** | | |
| pIMAY | Temperature-sensitive allelic exchange plasmid for staphylococci | Simon Heilbronner [7] |
| pIMAY-Δ*lip1* | pIMAY carrying the *lip1*(*gehA*) deletion construct. | This study |
| pIMAY-Δ*lip2* | pIMAY carrying the *lip2* (*gehB*) deletion construct. | This study |
| pEmpty (pALC2073) | *E. coli*-*S. aureus* shuttle vector; contains Pxyl/tet | David E. Heinrichs [9] |
| p*lip1* | pALC2073 containing the *lip1* coding region | This study |
| p*lip2* | pALC2073 containing the *lip2* coding region | David E. Heinrichs [3] |
| p*lip2*$^{S412A}$ | pALC2073 containing the mutated *lip2* coding region to generate Lip2 S412A | This study |
| pET28a(+)-*lip2* | pET28a(+) for Lip2 expression | David E. Heinrichs [3] |
| pET28a(+)-*lip2*$^{S412A}$ | pET28a(+) for Lip2 S412A expression | David E. Heinrichs [3] |

### Supplemental references


1. Duthie ES, Lorenz LL (1952) Staphylococcal coagulase; mode of action and antigenicity. *J Gen Microbiol* **6:** 95-107

2. Horsburgh MJ, Aish JL, White IJ, Shaw L, Lithgow JK, Foster SJ (2002) sigmaB modulates virulence determinant expression and stress resistance: characterization of a functional rsbU strain derived from Staphylococcus aureus 8325-4. *J Bacteriol* **184:** 5457-5467

3. Cadieux B, Vijayakumaran V, Bernards MA, McGavin MJ, Heinrichs DE (2014) Role of lipase from community-associated methicillin-resistant Staphylococcus aureus strain USA300 in hydrolyzing triglycerides into growth-inhibitory free fatty acids. *J Bacteriol* **196:** 4044-4056

4. Fey PD, Endres JL, Yajjala VK, Widhelm TJ, Boissy RJ, Bose JL, Bayles KW (2013) A genetic resource for rapid and comprehensive phenotype screening of nonessential Staphylococcus aureus genes. *mBio* **4:** e00537-00512

5. Nguyen MT, Luqman A, Bitschar K, Hertlein T, Dick J, Ohlsen K, Bröker B, Schittek B, Götz F (2018) Staphylococcal (phospho)lipases promote biofilm formation and host cell invasion. *Int J Med Microbiol* **308:** 653-663

6. Wang R, Braughton KR, Kretschmer D, Bach TH, Queck SY, Li M, Kennedy AD, Dorward DW, Klebanoff SJ, Peschel A*, et al.* (2007) Identification of novel cytolytic peptides as key virulence determinants for community-associated MRSA. *Nat Med* **13:** 1510-1514

7. Monk IR, Shah IM, Xu M, Tan MW, Foster TJ (2012) Transforming the untransformable: application of direct transformation to manipulate genetically Staphylococcus aureus and Staphylococcus epidermidis. *mBio* **3:**

8. Monk IR, Tree JJ, Howden BP, Stinear TP, Foster TJ (2015) Complete Bypass of Restriction Systems for Major Staphylococcus aureus Lineages. *mBio* **6:** e00308-00315

9. Bateman BT, Donegan NP, Jarry TM, Palma M, Cheung AL (2001) Evaluation of a tetracycline-inducible promoter in Staphylococcus aureus in vitro and in vivo and its application in demonstrating the role of sigB in microcolony formation. *Infect Immun* **69:** 7851-7857
