## Supplemental Table 3 for "Lipase-mediated detoxification of host-derived antimicrobial fatty acids by *Staphylococcus aureus*"

**Table S3. Primers.**

| Oligonucleotide | Sequence (5' – 3') |
|---|---|
| For deletion of *lip1* (pIMAY-Δ*lip1*): | |
| KpnI-*lip1* ups-fwd | CTAGAAGGTACCGGTGTCGGCATGATATTGCG |
| *lip1* ups-rev | CAAGCATAATTTATAAAGTAAAGGGAGG |
| SOEING-*lip1* dwn-fwd | CCTCCCTTTACTTTATAAATTATGCTTGACTTTTCATCATTGTCAGCACCTC |
| SacI-*lip1* dwns-rev | GCATCCGAGCTCGTAGGATACTTACTTTGAGGGAAG |
| For deletion of *lip2* (pIMAY-Δ*lip2*) | |
| KpnI-*lip2* ups-fwd | GTAGAGGTACCGTATGCCCACTAAACTATAGAC |
| *lip2* ups-rev | TCCTCTTAACATATAATCACCTC |
| SOEING-*lip2* dwn-fwd | GAGGTGATTATATGTTAAGAGGAGCAAGTTAAATTCATCTTCTG |
| SacI-*lip2* dwns-rev | CATGTTGAGCTCCGTTTATGCACGTGGCACAGG |
| For site-directed mutagenesis of p*lip2* into p*lip2*$^{S412A}$ | |
| *lip2*$^{S412A}$-fwd | GTAGGGCATGCTATGGGTGG |
| *lip2*$^{S412A}$-rev | CAAGATGTACCTTTTTACCAGG |
