## Supplemental Table 4 for "Lipase-mediated detoxification of host-derived antimicrobial fatty acids by *Staphylococcus aureus*"

**Table S4. MS/MS experiment of SWATH windows with m/z range, accumulation time (Acc. time) and collision energy (CE).**

| Experiment | Scan type | Acc. Time (ms) | ESI (+) Start (m/z) | ESI (+) Stop (m/z) | ESI (+) CE (V) | ESI (-) Start (m/z) | ESI (-) Stop (m/z) | ESI (-) CE (V) |
|---|---|---|---|---|---|---|---|---|
| 1 | MS Full Scan | 50 | 50.0 | 1250.0 | 10 | 50 | 1250.0 | -10 |
| 2 | SWATH | 31 | 50.0 | 217.6 | 45±15 | 50 | 213.5 | -45±15 |
| 3 | SWATH | 31 | 216.6 | 340.3 | 45±15 | 212.5 | 271.4 | -45±15 |
| 4 | SWATH | 31 | 339.3 | 441.4 | 45±15 | 270.4 | 314.6 | -45±15 |
| 5 | SWATH | 31 | 440.4 | 524.9 | 45±15 | 313.6 | 382.6 | -45±15 |
| 6 | SWATH | 31 | 523.9 | 571.6 | 45±15 | 381.6 | 427.5 | -45±15 |
| 7 | SWATH | 31 | 570.6 | 643.4 | 45±15 | 426.5 | 464.3 | -45±15 |
| 8 | SWATH | 31 | 642.4 | 687.3 | 45±15 | 463.3 | 501.0 | -45±15 |
| 9 | SWATH | 31 | 686.3 | 720.1 | 45±15 | 500.0 | 540.8 | -45±15 |
| 10 | SWATH | 31 | 719.1 | 740.1 | 45±15 | 539.8 | 617.5 | -45±15 |
| 11 | SWATH | 31 | 739.1 | 755.0 | 45±15 | 616.5 | 680.3 | -45±15 |
| 12 | SWATH | 31 | 754.0 | 764.1 | 45±15 | 679.3 | 697.1 | -45±15 |
| 13 | SWATH | 31 | 763.1 | 775.1 | 45±15 | 696.1 | 724.0 | -45±15 |
| 14 | SWATH | 31 | 774.1 | 786.1 | 45±15 | 723.0 | 749.0 | -45±15 |
| 15 | SWATH | 31 | 785.1 | 793.1 | 45±15 | 748.0 | 775.6 | -45±15 |
| 16 | SWATH | 31 | 792.1 | 806.1 | 45±15 | 774.6 | 793.1 | -45±15 |
| 17 | SWATH | 31 | 805.1 | 814.2 | 45±15 | 792.1 | 811.0 | -45±15 |
| 18 | SWATH | 31 | 813.2 | 829.6 | 45±15 | 810.0 | 832.6 | -45±15 |
| 19 | SWATH | 31 | 828.6 | 842.7 | 45±15 | 831.6 | 854.1 | -45±15 |
| 20 | SWATH | 31 | 841.7 | 903.3 | 45±15 | 853.1 | 861.2 | -45±15 |
| 21 | SWATH | 31 | 902.3 | 1250.0 | 45±15 | 860.2 | 1050.0 | -45±15 |
